# TAOK1 restrains cerebral cortex size and determines neural progenitor fate through YAP1 suppression

**DOI:** 10.64898/2026.11.27.769681

**Authors:** Sarah John, Yixuan Zhou, Daphnée Michelle Marciniak, Michael-Claude G Beltejar, Jennifer Hesson, Chris Cavanaugh, Moira Cornell, Dhriti Manjunath, Julie Mathieu, Mary Regier, Jessika Johannsen, Shao-En Ong, Smita Yadav

## Abstract

The size and complexity of the human cerebral cortex results from precisely coordinated neural progenitor expansion, differentiation, and fate trajectories for which the controlling molecular programs are not well understood^1,2^. Pathogenic missense and deletion variants in the protein kinase TAOK1 cause a neurodevelopmental disorder characterized by intellectual disability and macrocephaly^3–5^. We generated human cortical organoids harboring catalytically-inactive TAOK1 disease variant and utilized quantitative phosphoproteomics along with single-cell transcriptomics to uncover disease mechanisms. Catalytically-inactive TAOK1 increases neural progenitor proliferation, accelerates cortical organoid growth, and alters the diversity of cortical cell-types. Phosphoproteomic analyses revealed impaired Hippo signaling with reduced inhibitory phosphorylation of YAP1 concomitant with its sustained nuclear accumulation and activation of TEAD dependent transcription. Progenitors derived from a patient with heterozygous *TAOK1* deletion also accumulate nuclear YAP1, extending this finding to a distinct clinical disease-associated TAOK1 lesion. Single-cell transcriptomics demonstrate premature emergence of astrocytic and inhibitory neuronal populations in mutant organoids. Pharmacological inhibition of YAP-TEAD signaling reduced cyclin-A2 labeling and suppressed perturbed Hippo pathway targets. These findings identify TAOK1 as a molecular brake on human cortical overgrowth, implicate previously unrecognized role for Hippo dysfunction in macrocephaly, and reveal pathogenic consequences of accelerated progenitor cycling for both progenitor expansion and fate.

## MAIN

Altered growth trajectory of the cerebral cortex in humans is an early indicator of a wide range of neurodevelopmental deficits^6,7^. Aberrantly increased head circumference or macrocephaly is associated with genetically well-defined forms of neurodevelopmental disorders^8^ caused by pathogenic variants in phosphatase and tensin homolog (PTEN)^9^, PI3K-AKT-MTOR pathway^10^, CHD8^11^, and the RAS-MAPK pathway^12^. Deletion and duplication of the same genetic region in gene loci such as 16p11.2 and 22q11.2, among others, confers susceptibility to opposing effects on head size^13,14^, further demonstrating the genetic bases of brain size control. Cellular processes that contribute to the trajectory and extent of brain growth include progenitor pool expansion, timing and balance of differentiation with division, hypertrophy of neuronal arbors and glial development^7^. The molecular programs governing the cellular processes that ultimately safeguard human brain size and its developmental trajectory, remain largely unknown.

Pathogenic variants in the serine-threonine kinase encoding gene *TAOK1 (Thousand And One amino acid Kinase 1*) cause a syndromic neurodevelopmental disorder (NDD) with macrocephaly, developmental delay, intellectual disability, and autism^3–5,15–17^. Despite its disease association, the precise function of TAOK1 in brain development and how its dysfunction leads to neuropathology remains unknown. A majority of patients with TAOK1 pathogenic variants exhibit macrocephaly (head circumference mean +2.0 standard deviation), observed both in utero and postnatally^4,16^. Other structural malformations of the cerebral cortex including periventricular nodular heterotopia have also been reported in TAOK1 patients^5^.

TAOK1 belongs to the TAO family of evolutionarily conserved serine-threonine protein kinases represented by a single gene *Tao* in invertebrates, but three distinct genes *Taok1*, *Taok2* and *Taok3* in vertebrates^18^. Structurally, TAOK1 retains the evolutionary conserved domain architecture of the invertebrate Tao kinase. The N-terminal catalytic domain is highly conserved and shared among TAO kinases and their spliced isoforms^18^, while the C-terminal regulatory domain is divergent between TAOK1, TAOK2 and TAOK3 conferring functional specialization^19,20^. The TAO homolog in *C. elegans*, Kin18, regulates membrane contractility in embryos through RHO1 GTPase ^21^. In *Drosophila*, Tao mutants cause defects in border cell migration during oogenesis, mushroom body development^22^, epithelial tissue growth^23^, neuroblast division in ventral nerve cord^24^, and dendritic morphogenesis^25^. *Taok1* knockdown *in utero* in mice led to abnormal cortical neuron migration^15^, while a *Taok1* haploinsufficiency mouse model developed anxiety and autism-like behavior deficits^26^. We previously reported that neurodevelopmental disorder associated *TAOK1* variants in the kinase domain are catalytically deficient^19^. However, the molecular mechanisms through which neurodevelopmental disorder associated TAOK1 variants perturb early human brain development are unknown.

We developed human cortical organoid disease model of TAOK1 neurodevelopmental disorder and utilized quantitative phosphoproteomics and single-cell transcriptomics to uncover aberrant signaling and cellular mechanisms that lead to cortical overgrowth. TAOK1 pathogenic variant caused accelerated progenitor proliferation and increased cortical organoid size due to impaired Hippo signaling that converged upon aberrant nuclear accumulation of the transcriptional co-activator Yes associated protein 1 (YAP1). These findings were mirrored in TAOK1 deletion patient-derived neural progenitors indicating a converging disease mechanism of impaired kinase function shared by patients with TAOK1 deletions and missense variants. Single-cell transcriptomic analyses of organoids reveal precocious emergence of astrocytic lineage cells and cortical-derived inhibitory neurons in TAOK1 mutant organoids which was validated through immunocytochemistry. Aberrant YAP1-TEAD dependent gene expression signature is observed across distinct cell clusters in TAOK1 mutant organoids. Finally, pharmacological activation of TAOK1 or inhibition of YAP1 rescue progenitor proliferation and target gene expression. Our findings suggest that perturbed developmental programming drives macrocephaly in TAOK1-associated neurodevelopmental disorder and reveal a previously unrecognized role for Hippo signaling in human cortical size and development.

### TAOK1 catalytic activity controls expansion of neural progenitors

Macrocephaly is observed in 83% of individuals with TAOK1 neurodevelopmental disorder (TAOK1-NDD)^4^. Upon mapping the heterozygous missense TAOK1 mutations associated with macrocephaly, we found the mutations clustered within the kinase domain (Fig.1a, b), indicating an essential role of the catalytic function of TAOK1 in regulating human cortical growth. We selected one such macrocephaly associated variant p.Ser111Phe that we have previously established as catalytically inactive^19^. The patient carrying TAOK1 S111F variant presented with severe global developmental delay, macrocephaly, and muscular hypotonia^3^ (Extended Data Fig.1a). To assess how loss of TAOK1 catalytic activity contributes to TAOK1-NDD including macrocephaly, we used CRISPR-Cas9 mediated genome editing and generated iPSC lines harboring heterozygous (TAOK1^S111F het^) and homozygous (TAOK1^S111F hom^) TAOK1 S111F mutation, together with isogenic control (Ctrl^WT^) lines in the WTC11 genomic background (Extended Data Fig.1b). We confirmed the genotype, normal karyotype and pluripotency of individual isolated clones of the edited hiPSC lines Ctrl^WT^, TAOK1^S111F het^ and TAOK1^S111F hom^ by Sanger sequencing (Extended Data Fig.1c, d). Dorsal forebrain neural progenitor cells (NPCs) were differentiation from iPSCs using dual SMAD inhibition^27,28^ (Extended Data Fig.2a-c). We assessed the proliferative capacity of synchronized Ctrl^WT^, TAOK1^S111F het^, and TAOK1^S111F hom^ iPSC derived NPCs at 1 hr and 24 hr after addition of nutrients, followed by immunostaining for proliferation markers Ki-67 expressed in all phases of cell cycle except G_0_, and Cyclin A2 which is high in cells in the S and G2 phase. Both TAOK1^S111F het^ and TAOK1^S111F hom^ NPCs exhibited elevated number of cycling cells relative to control NPCs (Fig.1c-e), indicating TAOK1 loss of catalytic activity in NPCs promotes enhanced cell-cycle entry and progression through S/G2 phase.

**Figure 1.**
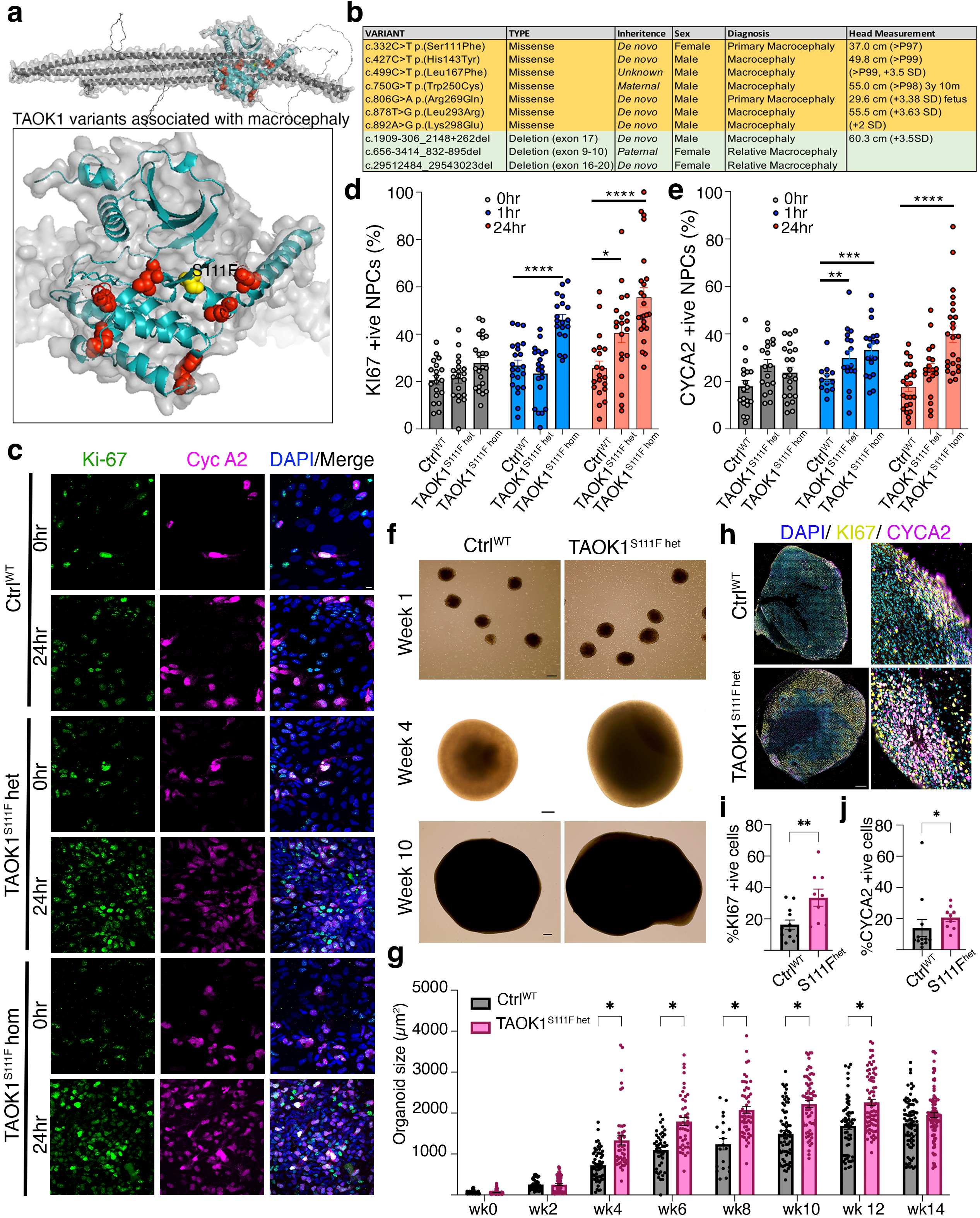
Neural progenitors and cortical organoids harboring catalytically inactive TAOK1 disease variant exhibit accelerated proliferation. (a) AlphaFold model of TAOK1 protein indicating location of missense variants (red) associated with macrocephaly. The variant S111F is highlighted in yellow. (b) List of TAOK1 missense variants and exon deletions associated with clinical macrocephaly. (c) Representative immunofluorescence images of wild-type (Ctrl^WT^), heterozygous (TAOK1^S111F het^), and homozygous (TAOK1^S111F hom^) iPSC derived neural progenitor cells (NPCs) in serum-starved condition (0 hr) and 24 h after serum readdition. Cells were stained for proliferation markers Ki-67 (green), Cyclin A2 (magenta), and DAPI (blue) counterstain. Scalebar, 10µm. (d) Percent Ki67-positive cells in Ctrl^WT^, TAOK1^S111F het^ and TAOK1^S111F hom^ NPC cultures at the indicated time points following re-addition of media is plotted. Data include independent experiments from three rounds of differentiation and are presented as mean ± SEM. *p =0.03, ****p<0.0001. (e) Proportion of CycA2-positive cells in Ctrl^WT^, TAOK1^S111F het^, and TAOK1^S111F hom^ NPC cultures at the indicated time points following re-addition of media is plotted. Data include independent experiments from three different rounds of differentiations and are presented as mean ± SEM. **p< 0.005, ***p< 0.001, ****p<0.0001. (f) Representative brightfield images of Ctrl^WT^ and TAOK1^S111F het^ organoids at weeks 1, 4, and 10 of cortical organoid differentiation. Scalebar, 100µm. (g) Bar chart plots the growth of Ctrl^WT^ and TAOK1^S111F het^ dorsal forebrain cortical organoids from week 0 to week 14 in vitro. Data include independent experiments from three different rounds of differentiations and are presented as mean area ± SEM. *p< 0.001. Significance was assessed by multiple Mann-Whitney test. (h) Representative images of day 100 cortical organoid sections stained for Ki-67 (yellow), Cyclin A2 (magenta) and DAPI (blue); inset (orange) is magnified in the right column. Scalebar, 100µm. (i) Percentage of Ki-67+ cells in day 100 Ctrl^WT^ and TAOK1^S111F het^ cortical organoid sections is plotted. Data are presented as means ± SEM (n = 4-6 organoids per condition). *p < 0.05; Student’s t test. (i) Percentage of CycA2+ cells in DIV 100 Ctrl^WT^ and TAOK1^S111F het^ cortical organoid sections is plotted. Data are presented as means ± SEM (n = 4-6 organoids per condition). *p <0.05; Student’s t test.

### Macrocephaly-like overgrowth phenotype in TAOK1 mutant cortical organoids

To determine the developmental consequences of accelerated NPC proliferation caused by inactivating TAOK1 mutation, we generated dorsal forebrain cortical organoids from control Ctrl^WT^ and TAOK1^S111F het^ hiPSCs using a directed differentiation protocol with SMAD and Wnt modulation^29^ (Extended Data Fig.2d, e). Ctrl^WT^ and TAOK1^S111F het^ hiPSCs generated spheroids with comparable overall size until 2 weeks. Neuroepithelial cells were organized in stereotypical ventricular zone–like architecture resembling the early developing cortex in both Ctrl^WT^ and TAOK1^S111F het^ organoids (Extended Data Fig.2e). By week 4, TAOK1^S111F het^ organoids exhibited a significant increase in size relative to controls (Fig.1f). The difference between the average sizes of organoids of mutant and the isogenic control continued to increase with overgrowth becoming most pronounced at week 8, when the maximum difference in mean organoid size (838.4μm^2^) between control and TAOK1 mutant organoids was observed (Fig.1f, g). Neuroprogenitor cells immunostained with Ki67+, CycA2+ were mostly localized to the apical surface of the ventricular zone (Extended Data Fig.2f). Consistent with organoid overgrowth, at week 8, TAOK1^S111F het^ organoids exhibited a significant increase in the proportion of proliferative cells, with elevated Ki67^+^ and cyclin A2^+^ progenitor population (Fig.1h-j). Subsequent growth of cortical organoids beyond week 12 decreased in both mutant and control organoids and was minimal beyond week 14. These data indicate that the catalytic activity of TAOK1 controls the developmentally timed expansion of the neural progenitor pool, perturbation of which leads to aberrant expansion associated with cortical overgrowth.

**Figure 2:**
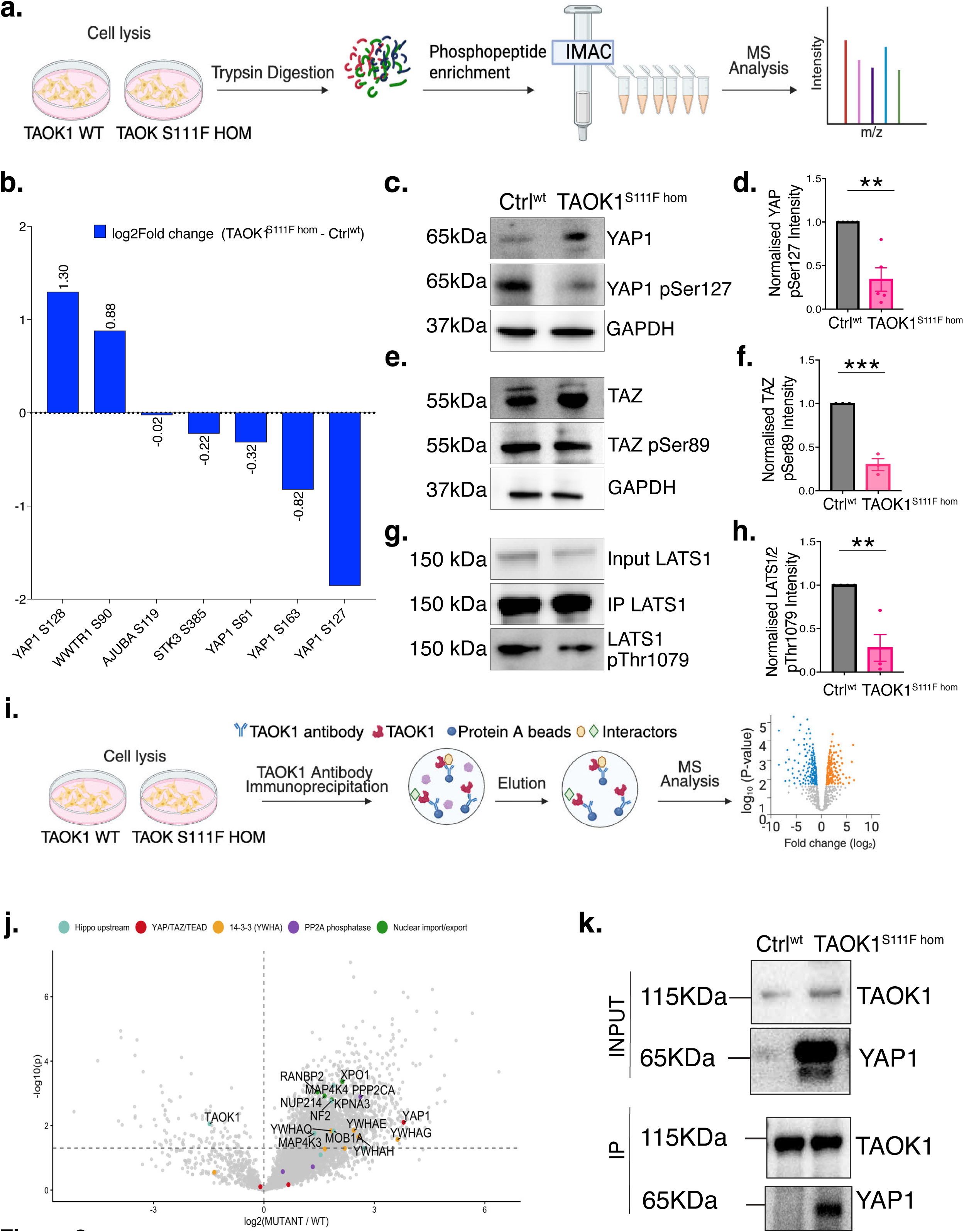
Hippo signaling dysfunction in TAOK1 kinase-dead mutant NPCs. **(a)** Workflow for the IMAC-based phosphopeptide enrichment and mass spectrometry analyses of phosphopeptides in WT and homozygous TAOK1 mutant NPCs. **(b)** Bar chart depicts foldchange in phosphopeptide intensity of key Hippo pathway regulatory sites identified in Ctrl^WT^ and TAOK1^S111F hom^ NPCs. Differential abundance was assessed using limma (*adj p = 0.01276). **(c)** Representative western blot of total YAP1 and phospho-YAP1 (Ser127) in lysates from Ctrl^WT^ and TAOK1^S111F hom^ NPCs. GAPDH level is used as loading control. **(d)** Normalized level of pS127 YAP1 to total YAP1 levels in Ctrl^WT^ and TAOK1^S111F hom^ NPCs is plotted. Normalized mean intensity ± SEM from five independent western blots. Statistical significance was assessed by two-tailed unpaired Student’s *t*-test (**p< 0.005, n=5). **(e)** Representative western blot of total TAZ/WWTR1 and phospho-TAZ (Ser89) in lysates from Ctrl^WT^ and TAOK1^S111F hom^ NPCs. GAPDH level is used as loading control. **(f)** Intensity of pS89 TAZ normalized to total TAZ levels in Ctrl^WT^ and TAOK1^S111F hom^ NPCs is plotted. Normalized mean intensity ± SEM from three independent western blots. Statistical significance was assessed by two-tailed unpaired Student’s *t*-test (***p =0.0005, n=3). **(g)** Endogenous LATS1/2 immunoprecipitated from Ctrl^WT^ and TAOK1^S111F hom^ NPC lysates, probed with total LATS1/2 and phospho-LATS1/2 (Thr1079) antibodies. **(h)** LATS1/2 Thr1079 phosphorylation levels normalized to total immumoprecipitated LATS1/2 in Ctrl^wt^ and TAOK1^S111F hom^ NPCs. Normalized mean intensity ± SEM from four independent western blots. Statistical significance was assessed by two-tailed unpaired Student’s *t*-test (**p <0.005, n=4). **(i)** Schematic depicts the immunoprecipitation–mass spectrometry (IP-MS) workflow used to identify TAOK1 interacting proteins from Ctrl^WT^ and TAOK1^S111F hom^ NPC lysates. **(j)** Volcano plot depicts proteins co-purifying with wildtype and S111F inactive TAOK1 mutant. Each point represents an identified protein, the x-axis shows log_2_ fold change in abundance, and the y-axis shows −log_10_(p-value). Dashed horizontal line indicates the significance threshold (p = 0.05, - log10(p) ∼ 1.30). Proteins are colored by functional category: Hippo pathway upstream regulators (blue), YAP/TAZ/TEAD (red), 14-3-3 proteins (yellow), PP2A phosphatase subunits (purple), and nuclear import/export proteins (green). Select proteins of interest are labeled by gene name. **(k)** Endogenous wildtype and mutant TAOK1 was immunoprecipitated from Ctrl^wt^ and TAOK1^S111F hom^ NPCs, respectively, and western blot shows immunoprecipitated proteins probed for TAOK1 and YAP1.

### Inactivation of Hippo pathway by catalytically deficient TAOK1

To isolate signaling abnormalities underlying hyperproliferation caused by the inactivating pathogenic TAOK1 variant, we performed label-free quantitative phosphoproteomics. Phosphoproteomes of Ctrl^WT^ and homozygous mutant TAOK1^S111F hom^ were compared as the latter completely lacks TAOK1 mediated phosphorylation events (Fig.2a). KEGG pathway analyses were performed on proteins differentially phosphorylated between Ctrl^WT^ and homozygous mutant TAOK1^S111F hom^ NPCs (Extended Data Fig.3a). Pathways including cell cycle (FDR = 3.9E-02 × 10⁻⁴, n = 8 genes), cellular senescence (FDR = 2.3E-03 × 10⁻⁴, n = 8 genes), and Hippo signaling (FDR = 3.9E-02 × 10⁻⁴, n = 8 genes) were identified suggesting that loss of TAOK1 catalytic activity impacts pathways governing NPC proliferative state. Unlike other growth pathways, dysfunction of Hippo signaling has not been associated with macrocephaly and the extent to which it plays a role human brain development is unknown.

**Figure 3:**
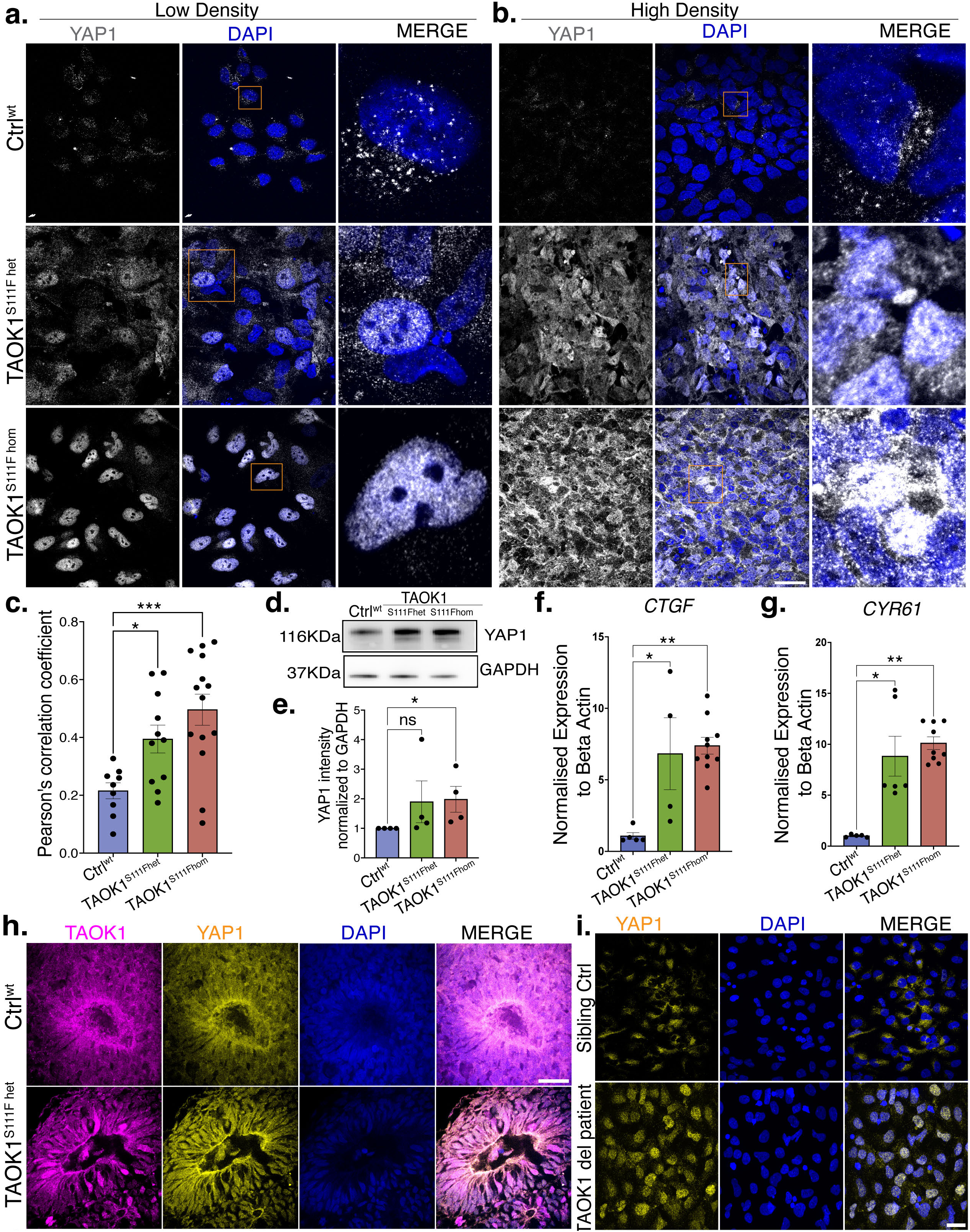
Aberrant nuclear YAP1 drives proliferation in TAOK1 mutant progenitors, cortical organoids and patient-derived samples. **(a)** Representative immunofluorescence images of low-density cultures of Ctrl^WT^, TAOK1^S111F het^, and TAOK1^S111F hom^ NPCs immunostained with YAP1 (gray) and DAPI (blue). Magnified inset (orange) is shown in right column. Scale bar, 25µm. **(b)** Representative immunofluorescence images of Ctrl^WT^, TAOK1^S111F het^, and TAOK1^S111F hom^ high-density NPC cultures immunostained for YAP1(gray) and DAPI (blue). Magnified inset (orange) is shown in right column. Scale bar, 25µm. **(c)** Pearson’s coefficient of colocalization between DAPI and YAP1 signals in Ctrl^WT^, TAOK1^S111F het^ and TAOK1^S111F hom^ NPCs is plotted. Statistical significance was assessed by Brown-Forsythe and Welch ANOVA test (* p< 0.05, *** p<0.0005). **(d)** Representative western blot shows total YAP1 in Ctrl^WT^, TAOK1^S111F het^ and TAOK1^S111F hom^ NPC lysates. GAPDH level is used as loading control. **(e)** Quantification of total YAP1 levels in Ctrl^WT^ and TAOK1^S111F hom^ NPCs from three independent western blots (*p< 0.05, n=4). **(f)** Expression of *CTGF* in Ctrl^WT^, TAOK1^S111F het^ and TAOK1^S111F hom^ NPCs as measured by qPCR. Significance was assessed by Kruskal–Wallis test (*p< 0.05, **p< 0.005). **(g)** Expression of *CYR61* in Ctrl^WT^, TAOK1^S111F het^ and TAOK1^S111F hom^ NPCs, measured by qPCR. Significance was assessed by Kruskal–Wallis test (*p<0.05, **p<0.005). **(h)** Representative confocal microscopy images of Ctrl^WT^ and TAOK1^S111F het^ dorsal forebrain cortical organoid sections immunostained for TAOK1 (magenta), YAP1 (yellow) and DAPI (blue). Scale bar, 25µm. **(i)** Representative immunofluorescence images of NPCs differentiated from patient iPSCs carrying a heterozygous TAOK1 deletion, immunostained for YAP1 (yellow) and DAPI (blue). NPCs differentiated from unaffected sibling control iPSCs serve as the control. Scale bar, 25µm

The role of Hippo pathway in *D. melanogaster* ventral nerve cord and neuroblast quiescence^24^ together with our findings of cortical overgrowth phenotype in TAOK1 mutant organoids, warranted further examination of the potential role of Hippo signaling in brain size control. The conserved Hippo pathway keeps proliferation in check through inhibitory phosphorylation of transcriptional co-activator YAP1 at Ser127 by LATS1/2 kinase, which leads to YAP1 degradation, thereby halting YAP-TEAD dependent transcriptional activity^30^. To further identify the proteomic signature of Hippo signaling dysfunction caused by TAOK1 mutation, we extracted all differentially phosphorylated sites attributed to known Hippo pathway components (Extended Data Fig.3b, c) and then restricted the analysis to those phospho-sites that have known regulatory functions (Fig.2b). We found that intensity of pSer127 of YAP1, the phosphorylation dependent 14-3-3 binding site on YAP1 which causes its cytoplasmic sequestration and degradation^31^, was reduced in TAOK1 mutant NPCs. Two LATS-dependent phosphorylation sites on YAP1^32^, pSer61 and pSer163, were similarly reduced in TAOK1 mutant NPCs. Conversely, phosphorylation of YAP1 at Ser128 and TAZ at Ser90 which antagonizes 14-3-3 binding and favors nuclear accumulation^33,34^ of YAP and TAZ, was increased in inactive TAOK1 mutant NPCs (Fig.2b). Consistent with the mass spectrometry data, reduced phosphorylation of YAP1 at Ser127, TAZ at Ser89, and LATS1 at Thr1079 were detected by Western blot in TAOK1 mutant lines (Fig.2c-h). However, through *in vitro* kinase assays, we did not observe direct phosphorylation of YAP1 by TAOK1 (Extended Data Fig.3d), indicating indirect regulation of YAP1 pS127 levels in TAOK1 mutant NPCs.

Since, YAP1 is not directly phosphorylated by TAOK1, and yet the effect on YAP1 is dependent on TAOK1 catalytic activity, we next tested whether TAOK1 associates with components of the Hippo signaling pathway in a catalytic activity dependent manner. Using an antibody against TAOK1, we affinity purified endogenous wildtype TAOK1 and kinase-dead TAOK1 S111F from Ctrl^WT^ and TAOK1^S111F hom^ mutant NPCs, respectively, and then used mass spectrometry to identify TAOK1 interactome in its active and kinase-dead states (Fig.2i). Notably, inactive TAOK1 showed increased association with core Hippo kinases and phosphatases, Hippo scaffolding proteins, 14-3-3 isoforms, and components of the nuclear import–export machinery (Fig.2j and Extended Data Fig.3e). One of the significant interactors of kinase-dead TAOK1 was YAP1. Co-immunoprecipitation confirmed the association between YAP1 and TAOK1 which was greatly potentiated by its catalytically off-state (Fig.2k). Since YAP1 drives proliferation through TEAD mediated gene transcription, and its phosphorylation levels are reduced in TAOK1 mutant NPCs, we further investigated whether its dysfunction contributes to hyperproliferation in TAOK1 mutant progenitors.

### Nuclear YAP1 accumulation and TEAD-dependent proliferation in TAOK1 mutant neural progenitors

Hippo signaling drives YAP1 phosphorylation and degradation which prevents nuclear transport and subsequent association of YAP1 with TEAD transcription factors, thereby suppressing gene programs that promote proliferation during organ growth^31^. In mammalian cells at high cell density, Hippo signaling turns on to degrade YAP1 and limit cellular growth^31,35^. Therefore, we used high cellular density as a paradigm to test whether Hippo signaling switch is impacted by TAOK1 mutation. In Ctrl^WT^ NPCs, as expected, YAP1 levels were low and entirely excluded from the nucleus. In contrast, mutant TAOK1^S111F het^ and TAOK1^S111F hom^ NPCs had abnormally elevated nuclear YAP1 even at high density (Fig.3a-c), indicating dysregulation of Hippo activation in response to external cues that promote growth inhibition. Consistent with the loss of pSer127 phosphorylation in TAOK1 mutant NPCs, elevated total levels of YAP1 protein were present in TAOK1^S111F het^ and TAOK1^S111F hom^ mutant NPCs compared to control NPCs (Fig.3d, e). Further, TAOK1 mutant NPCs had elevated mRNA levels of the canonical YAP-TEAD targets *CTGF* and *CYR61* that promote cellular proliferation (Fig.3f, g), indicating functional activation of nuclear YAP1. To test whether YAP1 nuclear-cytoplasmic shuttling was also perturbed during cortical development, we immunostained week 4 old Ctrl^WT^ and TAOK1^S111F het^ cortical organoid sections, and found elevated level of nuclear YAP1 within the proliferative zone in mutant organoids but not in control (Fig.3h). Along with the nuclear distribution of YAP1, we observed nuclear localization of TAOK1 in S111F mutant cells but not in control. The co-distribution of kinase-dead mutant TAOK1 with YAP1 in the nucleus is congruent with increased TAOK1-YAP1 association that we observe in kinase-dead state in our IP-MS experiment. Next, we determined if aberrant nuclear accumulation of YAP1 would be convergent phenotype and present broadly in patients with TAOK1 variants. We reprogrammed iPSCs from peripheral blood monocytes obtained from a 7yr old female with cognitive development and speech delay diagnosed with a heterozygous TAOK1 deletion (TAOK1-del) alongside their unaffected siblings (Extended Data Fig.4a-d). Differentiated NPCs from both patient and sibling controls showed normal expression of FOXG1 and PAX6 (Extended Data Fig.4e). When tested for YAP1 localization, however, TAOK1-del patient NPCs showed both increased level of total YAP1 and enhanced nuclear YAP1 (Extended Data Fig.4f). In contrast, YAP1 was primarily detected in the cytoplasm in the two sibling derived NPCs (Fig.3i and Extended Data Fig.4f). These data extend our findings of aberrant nuclear YAP1 as a biomarker of TAOK1 neurodevelopmental disorder across both kinase-dead TAOK1 missense variants and variants that lead to TAOK1 haploinsufficiency.

**Figure 4:**
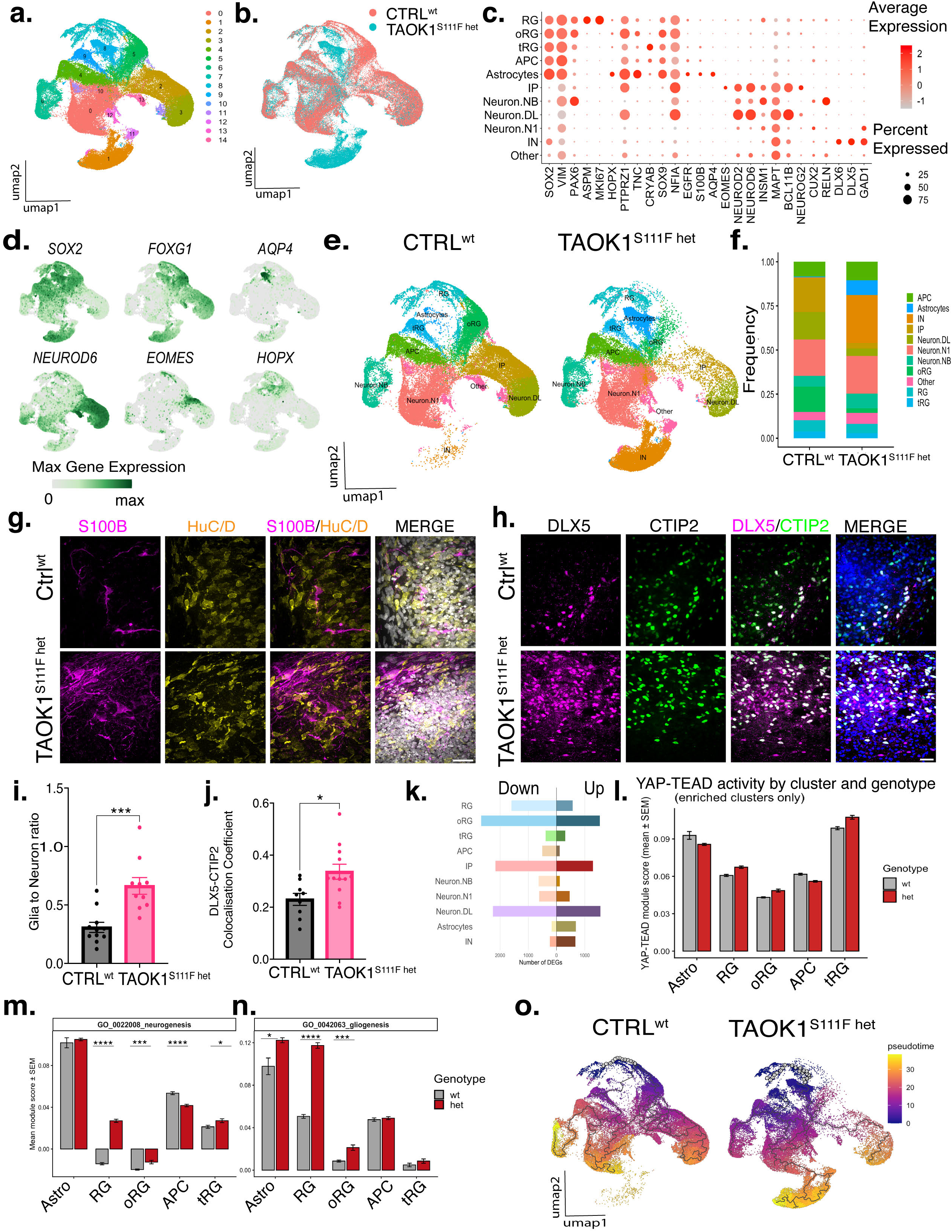
Transcriptomic analyses reveal altered cell fate toward astrocyte and interneuron lineages in TAOK1 mutant organoids. **(a)** UMAP plots of scRNA-seq data acquired from Ctrl^wt^ and TAOK1^S111F het^ dorsal forebrain cortical organoids at day 100, with cells grouped by unsupervised clustering and **(b)** by genotype. **(c)** Dot plot shows marker gene expression across annotated cell types. Color intensity reflects expression level (dark red, high expression); dot size reflects the percentage of cells expressing each marker. **(d)** Feature plots of key marker genes confirming cell identity: progenitors (*SOX2*), intermediate progenitors (*EOMES*), outer radial glia (*HOPX*), excitatory neurons (*NEUROD6*), astrocytes (*AQP4*), and dorsal telencephalic lineage (*FOXG1*). **(e)** UMAP plot showing cell-type annotations in Ctrl^WT^ and TAOK1^S111F het^ cortical organoids by cluster. **(f)** Cell-type proportions in Ctrl^WT^ and TAOK1^S111F het^ cortical organoids stratified by genotype. **(g)** Representative immunofluorescence images of S100B and HuC/D staining in Ctrl^WT^ and TAOK1^S111F het^ cortical organoid sections at DIV 100. Scale bar, 25µm. **(h)** Representative immunofluorescence images of DIV 100 Ctrl^WT^ and TAOK1^S111F het^ cortical organoid sections immunostained for CTIP2 and DLX5 to label inhibitory neurons. Scale bar, 25µm. **(i)** Bar plot quantifies (mean ± SEM) the glia-to-neuron ratio, calculated as the ratio of S100B+ to HuC/D+ cells per ROI in Ctrl^WT^ and TAOK1^S111F het^ cortical organoid sections at DIV 100 (n=6 organoids per genotype). Statistical significance was assessed by Mann–Whitney test, ***p≤ 0.001. **(j)** Bar plot depicts the Pearson correlation coefficient (mean ± SEM) for colocalization between CTIP2+ and DLX5+ nuclei in Ctrl^WT^ and TAOK1^S111F het^ cortical organoid sections at DIV 100 (n=6 organoids per genotype). Significance was assessed by Mann–Whitney test, *p < 0.05. **(k)** Number of DEGs between Ctrl^WT^ and TAOK1^S111F het^ cortical organoids in each cell cluster of the scRNA-seq dataset. **(l)** YAP–TEAD activity score across cell clusters in Ctrl^wt^ and TAOK1^S111F het^ cortical organoids, calculated from a (164 gene) YAP–TEAD target signature. **(m)** YAP–TEAD activity score in the clusters enriched in **(l)**, calculated using only neurogenesis-associated genes from the signature. Bars show the mean activity score per cluster. RG: adj p= 9.31E-102, oRG: adj p= 6.15E-04, tRG: adj p= 2.83E-09, APC: adj p= 9.31E-102. **(n)** YAP–TEAD activity score in the clusters enriched in **(l),** calculated using only gliogenesis-associated genes from the signature. Bars show the mean activity score per cluster. RG: adj p= 5.36E-85, oRG: adj p= 1.76E-04, Astrocytes: adj p= 4.45E-02. **(o)** UMAP embeddings of cells from Ctrl^WT^ and TAOK1^S111F het^ cortical organoids colored by pseudotime, with the trajectory inferred using Monocle 3. The root was set in the radial glia (RG) population (black-outlined cells; pseudotime = 0). Pseudotime trajectories progress from purple (early) to yellow (late).

### Loss of TAOK1 activity shifts cell fate from excitatory to glial and inhibitory lineages

Unlike mice where YAP1 is mostly restricted to apical radial glia (RG), in gyrencephalic species, it is widely expressed within the subventricular zone including outer radial glia (oRG) and intermediate progenitors (IPs) in the brain^36^. YAP1 signaling regulates the balance of stem cell renewal, differentiation, and quiescence^37^. To determine how aberrant YAP1 activation caused by loss of TAOK1 activity might alter cortical development, we performed single-cell RNA sequencing (scRNA-seq) of week 14 TAOK1^S111F het^ mutant and isogenic Ctrl^WT^ cortical organoids. After quality control, 88,533 cells were retained: 45,669 WT, 42,864 HET; median 3,142 genes and 6,847 UMIs per cell (Extended Data Fig.5a-b). Unsupervised clustering resolved 14 transcriptionally distinct populations (Fig.4a, b and Extended Data Fig. 5c). Using canonical marker genes (Fig.4c, d), clusters were assigned to the major cortical cell types: radial glia *(VIM, PAX6, SOX2, MKI67*), outer radial glia *(HOPX, PTPRZ1*), truncated radial glia *(CRYAB, LIFR, EGFR*), neurons *(MAPT, BCL11B, NEUROD2, NEUROD6*), astrocytes/astroglia *(SOX9, NFIA, S100B, AQP4, GFAP)*, and GABAergic inhibitory neurons *(DLX5, DLX6, GAD1*). These cell-type assignments (Extended Data Fig.5f) corelated well with published single-cell meta-atlas of the developing human cortex^38^. Analysis of cell-type proportions show that although cells from TAOK1^S111F het^ mutant organoids mapped onto the same cell types as controls, their distribution was heavily skewed toward astrocytes and inhibitory neurons at week 14 (Fig.4e, f and Extended Data Fig.5d, e). Specifically, TAOK1^S111F het^ organoids generated significantly greater number of astrocytic progenitors (*SOX9, SPARCL, TNC, VIM*) and mature astrocytes (*AQP4, S100B, SLC1A3*) in comparison with controls (Fig.4e and Extended Data Fig.5e). Consistent with this, immunostaining showed an increased ratio of S100B⁺ and SOX9⁺ glial cells to HUC/D^+^ neurons in TAOK1 mutant organoids (Fig. 4g, i and Extended Data Fig.6a, b). Within the astrocyte cluster of TAOK1 mutant organoids, astrocyte-maturation genes (*AQP4, S100B, BCAN,* and *APOE*) were highly expressed, whereas pan-reactive and neurotoxic markers^39^ (*SERPINA3, LCN2, CCL2,* and *CD44*) were minimally expressed or absent (Extended Data Fig.6c, d), indicating precocious maturation of the glial lineage rather than reactive gliosis. Similarly, immunostaining showed a significant increase in the number of CTIP2⁺/DLX5⁺ inhibitory neurons in the TAOK1 mutant organoids compared to controls (Fig.4h and Extended Data Fig.5d). DLX5⁺ neurons also expressing CTIP2⁺ were abundantly present in TAOK1 mutant neurons while very few DLX5+ neurons were present in control organoids (Fig.4j). Further analyses of the CTIP2⁺/DLX5⁺ neuron population revealed six transcriptionally distinct subclusters (Extended Data Fig.6e-g). The majority of subclusters (0, 2, 3, 4 and 6) expressed the lateral ganglionic eminence (LGE) marker *MEIS2*^40^ and retained progenitor marker like *SOX2*, *HES1* and *DLX2*, suggesting these represent transitional interneuron progenitor states rather than terminally specified subtypes. Notably, a distinct subcluster 5 showed a caudal ganglionic eminence (CGE) derived signature, marked by high *ADARB2*, *SCGN* and *CALB2* expression similar to the transcriptional identity of cortical born interneurons^41^. Cells in subcluster 1 co-expressed a subset of CGE marker *ADARB2* and medial ganglionic eminence (MGE) marker *LHX6*, indicating a mixed or intermediate identity. This reciprocal reorganization of cell fate, in which the excitatory lineage contracts as inhibitory and glial lineages expand reflects a broader shift in the output of the dorsal progenitors in mutant TAOK1 cortical organoids.

**Figure 5:**
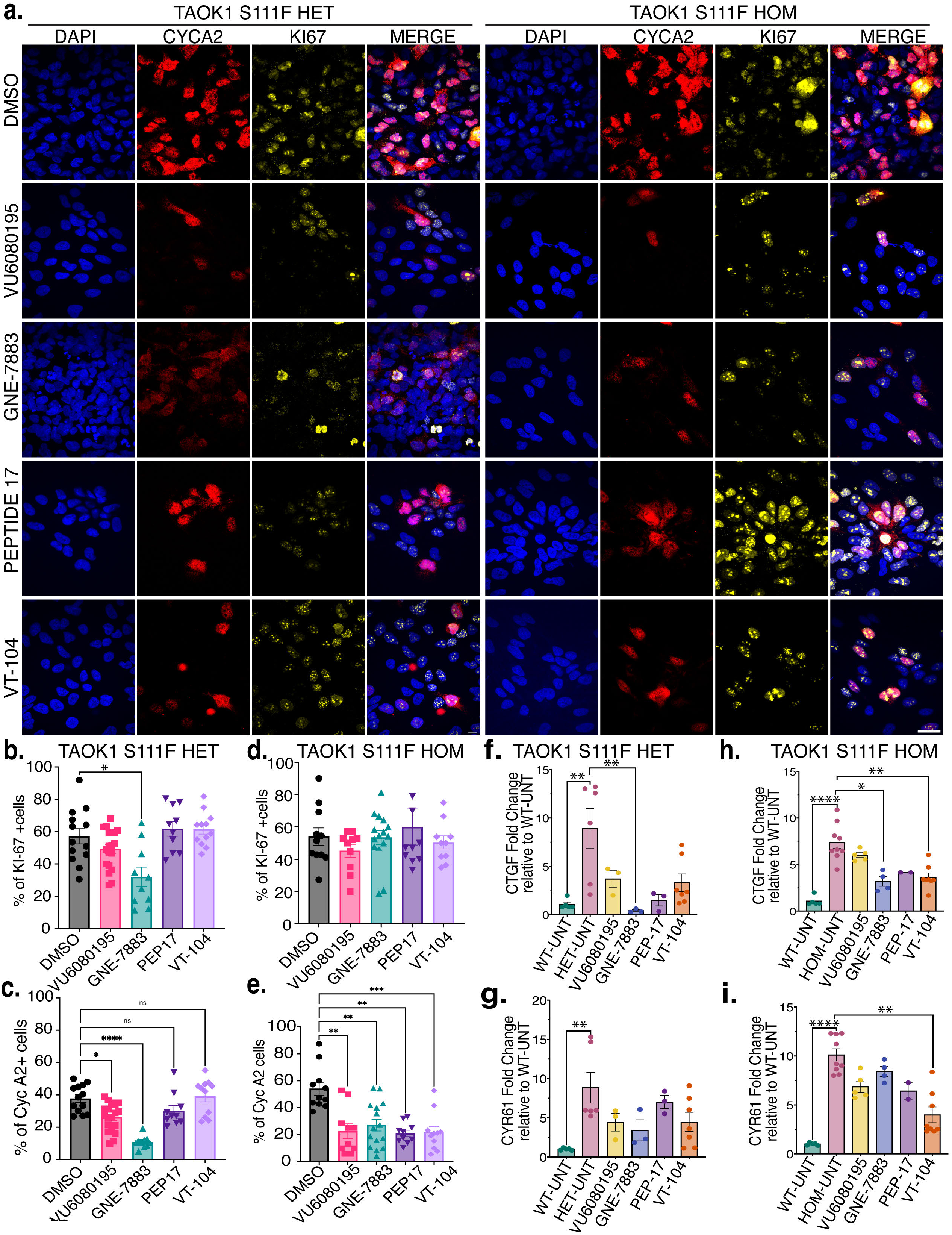
Targeting the TAOK1–YAP1–TEAD axis reverses excessive proliferation in TAOK1 mutant neural progenitors. **(a)** Representative immunofluorescence images of TAOK1^S111F het^ and TAOK1^S111F hom^ NPCs 24 h after treatment with DMSO, TAO activator VU6080195 (300nM), or YAP-TEAD inhibitors GNE-7883 (100nM), PEPTIDE-17 (100nM), and VT-104 (100nM). NPCs were immunostained for proliferation markers Ki-67 (yellow) and Cyclin A2 (red). Scale bar, 25µm. **(b)** Percentage of Ki-67⁺ cells in TAOK1^S111F het^ NPCs 24 h after treatment with DMSO, VU6080195, GNE-7883, PEPTIDE-17, or VT-104 is plotted. Data are presented as mean ± SEM from three independent differentiations. Significance was assessed by ANOVA, Kruskal–Wallis test (*p < 0.05). **(c)** Percentage of Cyclin A2⁺ cells in TAOK1^S111F het^ NPCs 24 h after treatment with DMSO, VU6080195, GNE-7883, PEPTIDE-17, or VT-104 is plotted. Data are presented as mean ± SEM from three independent differentiations. Significance was assessed by ANOVA, Kruskal–Wallis test (*p < 0.05, ****p ≤ 0.0001). **(d)** Percentage of Ki-67⁺ cells in TAOK1^S111F hom^ NPCs 24 h after treatment with DMSO, VU6080195, GNE-7883, PEPTIDE-17, or VT-104 is plotted. Data are presented as mean ± SEM from three independent differentiation. Significance was assessed by Kruskal–Wallis test. **(e)** Quantification of the percentage of Cyclin A2⁺ cells in TAOK1^S111F hom^ NPCs 24 h after treatment with DMSO, VU6080195, GNE-7883, PEPTIDE-17, or VT-104. Data are presented as mean ± SEM from three independent differentiations; significance was assessed by ANOVA, Kruskal–Wallis test (**p ≤ 0.01, ***p ≤ 0.001). **(f)** Bar graph of relative mRNA levels of *CTGF* and **(g)** *CYR61* in TAOK1^S111F het^ NPCs 24 h post-drug treatment, measured by qPCR. Significance was assessed by ANOVA, Kruskal–Wallis test (*p≤ 0.05, **p≤ 0.01). **(h)** Bar graph of relative mRNA levels of *CTGF* and **(i)** *CYR61* in TAOK1^S111F hom^ NPCs 24 h post-drug treatment, measured by qPCR. Data are presented as mean ± SEM from three independent differentiations; significance was assessed by ANOVA, Kruskal–Wallis test (* p≤ 0.05, **p ≤ 0.01, ***p ≤ 0.0001).

To determine which cells during cortical development were most transcriptionally affected by TAOK1 pathogenic variant, we quantified the number of differentially enriched genes (DEGs) in a cell cluster dependent fashion. The largest number of DEGs between TAOK1 mutant and control organoids were detected in RG, oRG, IP and deep layer neurons (DLN) population (Fig.4k). Since these populations also robustly express YAP1, we next assessed YAP-TEAD transcriptional activity using a curated signature of 164 distinct YAP-TEAD target genes previously reported to be expressed during cortical development^42^. Each cell was scored for YAP-TEAD activity using Seurat’s module score function which reports the average expression of the gene set relative to expression-matched control genes (Fig.4l). Astrocytes, RG, oRG, astrocyte progenitor cells (APCs), and truncated radial glia (tRG) showed enriched YAP-TEAD signature in TAOK1 mutant organoids compared to control (Fig.4l). Further, to identify the biological programs impacted by aberrant increase in YAP-TEAD activity in TAOK1 mutant organoids, we partitioned the YAP-TEAD signature by Gene Ontology (GO) annotation. TEAD genes annotated to neurogenesis (GO:0022008) and gliogenesis (GO:0042063) modules were used to build two distinct YAP-TEAD signature modules. The YAP-TEAD dependent neurogenesis module scored significantly higher in TAOK1 mutant RG than in control RG (Fig.4m). Gliogenesis module was more broadly impacted by kinase-dead TAOK1 mutation and YAP-TEAD gliogenesis genes were significantly higher in TAOK1 mutant RG, oRG, and astrocytes (Fig.4n). Among YAP-TEAD dependent neurogenic targets, *NTRK2* and *ASCL1* were significantly upregulated in mutant RG relative to control (Extended Data Fig.7a). Notably, gliogenic YAP-TEAD target genes *EGFR* and *METRN* were significantly upregulated in TAOK1 mutant RG compared to control (Extended Data Fig.7b). In astrocytes, we also observed significantly increased expression of YAP-TEAD effector gene *CCN1* (*CYR61*) (Extended Data Fig.7a, b), which has been reported to promote the maturation of inhibitory neurons^43^. Pseudotime reconstruction of developmental trajectories in control organoids revealed a progression from radial glial cells through intermediate progenitor and outer radial glial (oRG) states before bifurcating toward neuronal and glial branches, consistent with the known temporal order of cortical differentiation (Fig.4o). Comparing control and TAOK1 mutant organoids along this shared trajectory, the TAOK1 mutant UMAP shows a visibly greater proportion of cells occupying high-pseudotime (yellow) states in the lower branch corresponding to the inhibitory neuron lineage relative to WT, where advanced pseudotime values are more restricted to a terminal excitatory deep layer neuronal population. Together, these results suggest that aberrantly elevated YAP-TEAD activity within progenitor populations contribute to altered cell fate changes observed in TAOK1 mutant organoids such as expanded astrocytes and GABAergic inhibitory neuronal populations.

### Pharmacological suppression of YAP1–TEAD rescues hyperproliferation in TAOK1 mutant NPCs

We utilized two orthogonal pharmacological approaches to determine whether targeting the YAP1-TEAD pathway would suppress increased proliferation in TAOK1 mutant progenitors. Direct interruption of the YAP1-TEAD interface was achieved by three mechanistically distinct small molecules: an allosteric pan-TEAD inhibitor GNE7883^44^, an auto-palmitoylation inhibitor VT-104 that blocks YAP1 binding to TEAD1-4^45^, and the YAP1 derived inhibitory 17-mer disulfide-cyclized peptide, peptide-17^46^, that serves as a competitive inhibitor for TEAD binding. The second approach utilized a small molecule pan-TAO kinase activator, VU6080195^47^, to increase TAOK1 catalytic activity. We treated TAOK1^S111F het^, and TAOK1^S111F hom^ mutant NPCs with the YAP1-TEAD inhibitors, and TAOK1 activator and then assessed proliferation by immunostaining (Fig.5a). While total number of Ki-67 positive TAOK1 mutant cells were unchanged (Fig.5b, d), treatment with GNE-7883 produced the largest reduction in the number of actively cycling NPCs (Fig.5c, e). These data indicate that activating TAOK1 or suppressing YAP1 restrain S-phase entry or mitotic progression of TAOK1 mutant NPCs rather than driving their bulk exit into quiescence. Gene expression analyses of drug treated NPCs showed that GNE7883 reduced expression of YAP-TEAD effector *CCN2* (*CTGF*) expression in both TAOK1^S111F het^ and TAOK1^S111F hom^ NPCs significantly compared to the isogenic control (Fig.5f, h). All four compounds lowered *CCN*1 (CYR61) expression in TAOK1 S111F het NPCs to levels that are not significantly higher than the isogenic control (Fig. 5g), whereas in TAOK1^S111F hom^ NPCs only VT-104 deceased *CCN1* levels (Fig.5i). While TEAD-directed compounds rescued proliferation in TAOK1^S111F hom^ mutant NPCs, the TAOK1 activator VU6080195, as expected, did not have any effect on the homozygous TAOK1 mutant line due to lack of functional copy that can be activated.

In summary, these findings together with our data in organoids and patient derived NPCs, demonstrate that accelerated proliferation in TAOK1 mutant NPCs is driven by unchecked YAP1-TEAD activity and is pharmacologically reversible. Restoring TAOK1 catalytic activity or blocking downstream YAP1–TEAD interaction each normalizes proliferation and target-gene expression, uncovering the TAOK1–YAP/TEAD axis as a therapeutically actionable mechanistic node for TAOK1-NDDs.

## DISCUSSION

In this study, using human cortical organoids, we demonstrate that loss of TAOK1 kinase activity drives abnormal cortical growth accompanied by enhanced proliferation of dorsal forebrain neural progenitors, increased generation of cortical-derived inhibitory neurons, and premature gliogenesis. We show that altered YAP1-TEAD transcriptional activity due to aberrantly sustained nuclear accumulation of YAP1 leads to accelerated proliferation in TAOK1 mutant organoids. These changes are mirrored in TAOK1 deletion patient-derived neural progenitors but not in unaffected sibling samples. Our findings establish dysregulation of the Hippo signaling effector YAP1 as the pathological driver of macrocephaly and neurodevelopmental deficits in TAOK1 patients.

The transcriptional co-activator YAP1 has been extensively studied in the context of brain development. Mechanical force, cell-contacts, and growth factors serve as signals that restrict entry of YAP1 into the nucleus via phosphorylation and subsequent degradation. Nuclear YAP1, on the other hand, is consistently observed in progenitors during normal embryonic brain development as well as in pathogenic states such as gliomas, where it drives transcriptional programs of growth. Extensive and sustained YAP1 expression within subventricular zone in both oRGs and IPs cells of gyrencephalic species like ferrets and humans compared to rodents underscores the critical importance of YAP1 mediated neocortical expansion in evolution^36^. Further, pharmacological treatment of human cortical tissue by verteporfin that inhibits YAP1 association with TEAD resulted in decreased abundance of cycling oRGs and IPs^36^. This, together with previous studies in mice demonstrate that the proliferative effect of YAP1 in the brain is through TEAD transcription^36,48^. Our findings are congruent with this, and show that increased proliferation in TAOK1 mutant dorsal forebrain progenitors is mediated by YAP1-TEAD transcription and can be blocked by YAP-TEAD inhibitors.

Complex mechanisms likely underlie YAP1 activation upon TAOK1 loss of kinase activity or TAOK1 haploinsufficiency. In HEK293 cells, TAOK1 has been shown to directly phosphorylate MST1/2, however, knockdown of TAOK kinases or MST1/2 kinase in HEK293 cells is not sufficient to inhibit YAP1 phosphorylation^49^. However, knockdown of LATS1/2 led to abolished YAP1 phosphorylation and robust YAP1-TEAD activation. TAOK1 can directly phosphorylate and activate LATS1/2, which is turn mediated YAP1 phosphorylation at the critical Ser217^49^. In mice, deletion of Lats1/2 leads to accumulation of Yap/Taz in the nucleus concomitant with accelerated cell division of apical RG cells^50^. Consistent with these, we observed lower levels of active pT1079 LATS1 and pS127 YAP1 in TAOK1 mutant NPCs. We found that TAOK1 does not directly phosphorylate YAP1 and analyses of TAOK1 mutant NPCs suggests an important contribution of TAOK1 in phosphorylation of LATS1/2 kinases that cannot be compensated in NPCs by MST1/2 kinases. Moreover, our data unraveled a non-catalytic scaffolding function of TAOK1 that is executed in its kinase-dead state. We discovered that TAOK1 is associated with protein complexes enriched in Hippo signaling proteins in its inactive state. Indeed, kinase-dead TAOK1 showed significantly increased association with YAP1, 14-3-3 proteins, and nuclear import/export factors compared to active TAOK1. Consequently, we find that in NPCs, inactive mutant TAOK1 is distributed inside the nucleus along with YAP1. The significance of nuclear entry of inactive TAOK1 and its increased protein-protein interaction with components of the Hippo signaling pathway is not yet clear in this context, but may be important for protecting YAP1 from phosphorylation by other kinases. TAOK1 has been reported to localize to the cytoplasm and the plasma membrane^19^. Although studies have implicated it in nuclear functions including homologous recombination^51^, its presence inside the nucleus has not, to our knowledge, been described. We note here that TAOK1 carries a classical bipartite nuclear localization signal (KR-X_10–12_-KRRK) that fits the motif sequence predicted to interact with KPNA3^52^. It is plausible that TAOK1 in its catalytically off-state complexes with YAP1 to engage the binding site of importin proteins and enters the nucleus. We found that proliferative effect of aberrant YAP1 in TAOK1 mutant NPCs can be pharmacologically inhibited by disrupting YAP-TEAD interaction, where compound GNE-7883 had the strongest effect. A small-molecule TAOK1 activator partially normalized *CTGF* expression in NPCs but failed to restore *CYR61* possibly reflecting differential sensitivity of TEAD target genes to residual TAOK1 activity, or promoter-specific co-factor requirements that warrant further investigation. Given the broad developmental role of YAP1, targeting it seems an unlikely strategy for TAOK1 neurodevelopmental disoder, unless temporal and cell specific YAP1 engagement can be designed. Upregulating the unaffected copy of TAOK1 through antisense oligonucleotides or micro-RNAs to normalize YAP1 phosphorylation and prevent nuclear accumulation on the other hand, presents an on-target direct approach that could be explored.

Emergence of mature astrocytes and expansion of inhibitory neurons at the expense of intermediate progenitors and outer radial glial cells observed in TAOK1 mutant organoids indicates temporally accelerated cortical development. Altered cell fate with accelerated neuronal differentiation has been observed in human organoid models of several other neurodevelopmental disorders, to date. Haploinsufficiency of three autism linked genes, *KMT5B*, *ARID1B*, and *CHD8*, cause asynchronous development of GABAergic cortical neurons and deep layer excitatory neurons^53^. Dysregulated tempo of neurogenesis has been reported in organotypic slices of primary human cortices with trisomy 21, marked by the precocious development of *ADARB2*+ cortical inhibitory neurons ^54^. We propose that the fate shift caused by TAOK1 mutation reflects an accelerated differentiation timeline based on two observations: First, YAP-TEAD target genes involved in neurogenesis and gliogenesis, including *ASCL1*, *NTRK2*, *EGFR* and *METRN*, are upregulated in mutant radial glia, suggesting that there is an early switch that is turned on in progenitor cells driving the expansion of astrocytes and neuronal fate. Second, DLX5⁺/FOXG1⁺ neurons peak in abundance in six-month-old organoids in a published cortical organoid atlas^55^, suggesting that TAOK1 mutant organoids reach this state precociously^48^. We observe that TAOK1 mutant organoids produce greater number of inhibitory neurons as well as astrocytes with a concomitant decrease in intermediate progenitors and outer radial glial cells. As multipotent progenitors, oRGs in the human cortex are prolific producers of upper-layer excitatory neurons^56^, astrocytes and oligodendrocyte precursors^57^, and DLX5+/GAD2+ inhibitory neurons^58^. We observe altered YAP-TEAD signature in progenitor cells including IPs, tRG, and oRG due to loss of TAOK1 catalytic function. It is plausible that YAP1 activation biases oRG output away from the excitatory intermediate progenitor trajectory and toward inhibitory and glial fates, with premature differentiation depleting the progenitor pool. This parallels the finding that LIF signaling, which expands oRGs, reduces excitatory IPCs while increasing interneurons^58^. Future investigations into YAP1 dependent or independent mechanisms that contribute to developmental tempo and cell fates changes will provide deeper insight into potential therapeutic strategy for TAOK1 neurodevelopmental disorder, and more broadly, will help define the role of TAOK1-YAP1 signaling during human cortical development.

## METHODS

### CRISPR/Cas9-mediated Genome Editing of iPSCs

To introduce the S111F mutation in the TAOK1 gene, 1 × 10^6 WTC-11 human iPSCs were electroporated with a pre-assembled Cas9 ribonucleoprotein (RNP) complex containing Cas9 protein (0.3–0.6 μM; Sigma) and gRNA (1.5–3 μM; Synthego), together with a single-stranded DNA (ssDNA) donor template (2–4 μM; IDT). Electroporation was performed using the Amaxa Nucleofector system with the Human Stem Cell Kit 2 in the presence of a ROCK inhibitor. Following electroporation, cells were allowed to recover and expand, and individual colonies were manually picked and transferred to 96-well plates for clonal expansion. Genomic DNA was extracted using QuickExtract DNA Extraction Solution (Epicentre, QE09050). The targeted region was amplified by nested PCR using the following primers: forward primer, TGTATCATTGGCTTGACTTTATGCA; reverse primer, TGGTTTGAACAAGAGGAGCAGA. PCR products were purified using ExoSAP-IT (Thermo Fisher Scientific) and submitted for Sanger sequencing (Genewiz) to identify clones carrying the desired S111F mutation. The gRNA sequence was ATATTGTTTAGGATCTGCTT, and the ssDNA donor template sequence was ATATATTTTAATGCTTTTATAACTTTTCCATGTAGCTTGTAATGGAATATTGTTTAGGATTTGC TTCGGATTTACTAGAAGGTAAGTTCCCTTTGATTATTTTTTAAAAAGTATTCACAGA. Two independent clones were established for each genotype (Extended Data Fig.1c). Genomic integrity was assessed by karyotyping, which showed no chromosomal abnormalities (Extended Data Fig.1e).

### Reprogramming of Patient PBMC into iPSC

Blood samples from TAOK1 deletion patient and siblings were obtained with consent at the University Medical Center Hamburg-Eppendorf (Universitätsklinikum Hamburg-Eppendorf) under the supervision of Dr. J. Johanssen. Isolation of peripheral blood mononuclear cells (PBMC) were performed as follows: 20-30 ml heparinized peripheral blood were layered on top of 10-15 ml Lymphoprep™ (STEMCELL Technologies). Samples were centrifuged at 900 x g for 22 minutes at 20°C in a Megafuge 40R (Heraeus). Acceleration and deceleration were set to 4. The PBMC layer at the plasma:Lymphoprep™ interface was removed and retained. PBMC were washed twice with PBS (Gibco) and centrifuged at 300 x g for 5 minutes at 20°C. Cell concentrations were analyzed using a DxH 500 hematology analyzer (Beckman Coulter) and adjusted to 2×106/ml in freezing medium (50% RPMI 1640, Gibco; 25% FCS, Gibco; 10% DMSO, AppliChem). Aliquots of 2×106 PBMC each were subsequently frozen to −80°C at a controlled rate of ≈1°C/min. Isolated peripheral blood mononuclear cells (PBMC) were reprogrammed into iPSCs (at University of Washington) using the non-integrating CytoTune-iPS 2.0 Sendai Reprogramming Kit according to the manufacturer’s instructions (Thermo Fisher Scientific, Cat. No. A16517). Emerging colonies were established on irradiated CF1 mouse embryonic fibroblasts (MEFs) in DMEM/F-12 supplemented with 20% knockout serum replacement, nonessential amino acids (NEAA), sodium pyruvate, and penicillin–streptomycin (all from Invitrogen), 0.1 mM β-mercaptoethanol (Sigma-Aldrich), 0.1 µM sodium butyrate (Sigma-Aldrich, Cat. No. B5887), and 50 nM suberoylanilide hydroxamic acid (SAHA; Cayman Chemical, Cat. No. 10009929). Cells were cultured at 37°C in a humidified atmosphere containing 5% CO₂ and 5% O₂. Established colonies were initially passaged using dispase and subsequently adapted to feeder-free culture on Matrigel-coated plates in mTeSR Plus medium. Genomic integrity of isolated clones was assessed by karyotyping, which showed no chromosomal abnormalities.

### IPSC Culture and Neural Differentiation

All hiPSCs were cultured using mTeSR media (STEMCELL Technologies) containing 10 μM ROCK Inhibitor (Selleck Chemical). Cultures were passaged with 3ml Accutase (STEMCELL Technologies) per 10cm plate every 5-7 days. All iPSC lines were differentiated into neural progenitor cells (NPCs) using a previously published protocol (Deshpande et al., 2017). In brief, iPSCs were treated with dispase (1 U/mL), mechanically detached using a cell scraper, and resuspended in mTeSR medium (STEMCELL Technologies) supplemented with ROCK inhibitor (10 μM; Selleck Chemical). Cell suspensions were transferred to uncoated T25 flasks to promote embryoid body (EB) formation. After 24–48 h (designated day 0), EBs were allowed to settle by gravity and were transferred to new uncoated T25 flasks containing neural induction medium comprising DMEM/F12 (Gibco) supplemented with 1% N-2 (Gibco), 1% non-essential amino acids (Gibco), heparin (2 μg/mL), and 1% penicillin/streptomycin (Gibco). Neural induction was performed under dual SMAD inhibition using SB431542 (5 μM; PeproTech) and LDN193189 (0.25 μM; PeproTech). On day 3, EBs were plated onto Matrigel-coated (1:50 dilution; Corning) 6-well plates in fresh neural induction medium containing SB431542 and LDN193189. Thereafter, cultures were maintained with medium changes every other day using neural induction medium lacking SB431542 and LDN193189, as required, until day 11 to facilitate neural rosette formation. On day 11, neuroepithelial structures were mechanically lifted using a cell scraper and transferred to uncoated T25 flasks in neural induction medium to generate neurospheres. Neurospheres were maintained with neural induction medium replaced every other day, as required, until day 25. On day 25, neurospheres were plated onto Matrigel-coated plates in STEMdiff Neural Progenitor Medium, resulting in establishment of NPC cultures.

### Cerebral Organoid Culture

Cerebral organoids were generated from Ctrl^wt^ and TAOK1^S111F het^ hiPSCs using a previously reported protocol (Yoon et al 2019). All hiPSCs were maintained on vitronectin (VTN-N; Life Technologies, A14700) in Essential 8™ medium (Life Technologies, A1517001) and passaged at 80–90% confluency using 0.5 mM EDTA (Life Technologies, 15575; 7 min, room temperature). For spheroid formation, hiPSCs were dissociated with Accutase (Innovative Cell Technologies, AT-104; 7 min, 37 °C) and seeded into AggreWell™800 plates (STEMCELL Technologies, 34815) at ∼3 × 10^6^ cells per well in Essential 8™ + Y-27632 (10 μM; Selleckchem). After 24 h, spheroids were transferred to ultra-low attachment dishes (Corning, 3262) in Essential 6™ (Life Technologies, A1516401) with dual SMAD inhibition (dorsomorphin, 2.5 μM; Sigma-Aldrich, P5499; SB-431542, 10 μM; Tocris, 1614); and media was changed daily. On day 6, spheroids were transitioned to neural medium (Neurobasal A (Life Technologies, 10888), B-27 without vitamin A (Life Technologies, 12587), GlutaMAX (1:100; Life Technologies, 35050)). The neural media was supplemented with EGF (20 ng/mL; R&D Systems, 236-EG) and bFGF (20 ng/mL; R&D Systems, 233-FB) for 19 days and, then with BDNF (20 ng/mL; PeproTech, 450-02) and NT-3 (20 ng/mL; PeproTech, 450-03). From day 43, organoids were maintained in neural medium without growth factors with medium changes every 4 days. Multiple independent experiments were performed to generate cerebral organoids from Ctrl^wt^ (n = 5) TAOK1^S111F het^ (n = 4). Each experiment generated 15-20 individual organoids of each genotype.

### Immunohistochemistry

Cryosections were rinsed in PBS to remove residual OCT, then permeabilized and blocked in PBS containing 10% normal goat serum (NGS) and 0.3% Triton X-100 for 1 h at room temperature. Sections were incubated overnight at 4 °C with primary antibodies diluted in PBS containing 2% NGS and 0.1% Triton X-100. After washing in PBS, sections were incubated with appropriate fluorophore-conjugated secondary antibodies diluted in PBS containing 2% NGS and 0.1% Triton X-100 for 1 h at room temperature, followed by PBS washes. Primary antibodies used were: anti-Ki-67 (mouse; 1:500; Cell Signaling Technology, “9449), anti–cyclin A2 (rabbit; 1:500), anti-SATB2 (mouse; 1:50; Abcam, ab92446), anti-CTIP2 (rat; 1:300; Abcam, ab18465), anti-TBR1 (rabbit; 1:300; Abcam, ab31940), anti-TBR1 (chicken; 1:200; Millipore, ab2261), anti-GFAP (rat; 1:500; Thermo Fisher Scientific, 13-0300) and anti-SOX2 (rabbit; 1:200; Cell Signaling Technology, 3579).

### Phosphoproteomics

#### Sample preparation for mass spectrometry

Isogenic Ctrl^wt^ and TAOK1^S111F hom^ NPC pellets (5 replicates per genotype) were lysed in 8 M urea, 100 mM Tris-HCl (pH 8.0–8.5), 5 mM TCEP and 10 mM chloroacetamide. Lysates were sonicated in an ice-cooled cup-horn sonicator (100% amplitude, 30 s on/30 s off, 10 min), shaken at 37 °C for 30 min and cleared at maximum speed for 10 min at 4 °C. Proteins were precipitated with four volumes of ice-cold acetone and 5% TCA at −20 °C overnight, pelleted at 6,000 rcf, washed once in ice-cold acetone, resolubilised in fresh 8 M urea/100 mM Tris and quantified (Pierce 660 nm assay). Protein concentration was normalized across samples and were digested with endoproteinase Lys-C (1:100, w/w) at 4 M urea for 2 h at 37 °C, then with trypsin (1:100, w/w) at 2 M urea overnight at 37 °C, diluting with TEAB at each step. Digests were acidified to 2% formic acid. A 5% aliquot was reserved for the unenriched proteome (for global protein level normalization); the remainder was desalted on Waters Oasis 30 mg C18 cartridges. Phosphopeptides were enriched on Fe(III)-charged IMAC (ThermoScientific A32992) beads (20 µl 50% slurry per sample) for 1 h at 25 °C in 80% acetonitrile/0.1% TFA. Beads were washed four times in the same buffer, and phosphopeptides eluted three times in 150 µl 50 mM K₂HPO₄/NH₄OH (pH 10.0) into acid-neutralized tubes. Eluates were dried, desalted by C18 StageTip and resuspended in 0.1% TFA.

#### DD nano liquid chromatography (LC)-MS analysis

Peptide samples were separated on an EASY-nLC 1200 System (Thermo Fisher Scientific) using 20 cm long fused silica capillary columns (100 μm ID, laser pulled in-house with Sutter P-2000, Novato CA) packed with 3 μm 120 Å reversed phase C18 beads (Dr. Maisch, Ammerbuch, DE). MS data was collected with a Thermo Fisher Scientific Orbitrap Fusion Lumos Tribrid MS system. For phosphoproteomics experiments, the LC gradient was 90 min. long with 5–35% B at 300 nL/min. LC solvent A was 0.5% (v/v) aq. acetic acid and LC solvent B was 80% acetonitrile in 0.5% (v/v) acetic acid. A data-dependent analysis (DDA) MS acquisition method was applied with 60K resolution MS1 and 30K HCD fragmentation Orbitrap MS2 scans in a cycle-time data-dependent mode. Data .raw files were analyzed by MaxQuant/Andromeda version 1.6.17.0 using protein, peptide and site FDRs of 0.01 and a score minimum of 40 for modified peptides, 0 for unmodified peptides; delta score minimum of 17 for modified peptides, 0 for unmodified peptides. MS/MS spectra were searched against the UniProt human database (updated Feb 20th, 2020). MaxQuant search parameters: Variable modifications included Oxidation (M), Acetyl (Protein N-term) and Phospho (S/T/Y). Carbamidomethyl (C) was a fixed modification. Max. missed cleavages was 2, enzyme was Trypsin/P and max. charge was 7. The MaxQuant “match between runs” feature was enabled. The initial search tolerance for FTMS and HCD MS2 scans was 20 ppm. MaxQuant result tables including PhosphoSites(STY).txt, proteinGroups.txt, and evidence.txt were processed in R. Zero intensities were converted to missing values. Intensities were log2 transformed and each sample column was median-normalized by subtracting its own log2 median. Missing values were not imputed. Rows were tested only when at least two Ctrl^wt^ and at least two TAOK1^S111F hom^ values of 5 replicates were observed. From 5,384 phosphosites identified by mass spectrometry, we tested for differential phosphorylation between mutant and wild-type NPCs using limma^59^. KEGG pathway over-representation analysis was performed on the genes that were differentially phosphorylated between Ctrl WT and *TAOK1* S111F homozygous samples (adjusted *P* ≤ 0.05) using ShinyGO (v0.85).

### Immunoprecipitation and Kinase Assay

Ctrl^wt^ NPCs were lysed in RIPA lysis buffer (20 mM Tris HCL pH 7.5, 130mM NaCl, 2mM EDTA, 500mM NaF, 25% V/V NP-40,10% w/v Sodium deoxycholate, 20% w/v SDS) with 1x cOmplete protease inhibitor (Sigma). Lysate was then spun for 10 minutes at 15,000g at 4C to pellet out cell debris. Protein-A beads (Pierce^TM^, 20333) were equilibrated with 2x wash buffer, followed by pre-clearing of 1 mL cell lysate with 15 uL equilibrated beads for 30-60 min at 4C rotating. 1 ug of rabbit primary antibody (anti-TAOK1, Proteintech, 26250-1-AP) was conjugated to 30 uL of Protein A beads during this pre-clear step, followed by addition of pre-cleared lysate to the conjugated beads for immunoprecipitation. Proteins were immunoprecipitated for 16 hours at 4C with rotation. Following IP, samples were washed 2x with HKT (25 mM HEPES pH 7.2, 1 M NaCl, 1% Triton X-100, 1mM DTT), 1x with high salt HKT (25 mM HEPES pH 7.2, 1 M NaCl, 1% Triton X-100, 1mM DTT) for 10 minutes, 2x with HK buffer (25 mM HEPES pH 7.2, 1 M NaCl, 1mM DTT), and 2x with kinase buffer (20 mM Tris HCl pH 7.5, 10 mM MgCl2, 1 mM DTT, 1x cOmplete protease inhibitor). Following bead washes, kinase labeling reaction was performed for 45 minutes at 30C on a shaking heat block (Thermo) in 625 kinase buffer with 0.5mM ATP (Sigma) or ATPgS (Thermo) and 2x Halt Phosphatase inhibitor (Thermo). Following the kinase reaction, labeling was quenched by the addition of EDTA to a final of concentration of 50mM, and thiophosphates were then alkylated with p-nitrobenzyl mesylate (pNBM, Sigma) at a final concentration of 6.67mM for 45 minutes at room temperature. After alkylation, the reaction was quenched by the addition of 4X Lammeli-SDS sample 632 buffer (Thermo) with 200mM DTT.

### IP-MS Identification of Interactome

The LC gradient was 90 min. long with 6–45% B at 300 nL/min. LC solvent A was 0.5% (v/v) aq. acetic acid and LC solvent B was 80% acetonitrile in 0.5% (v/v) acetic acid. Data was collected using a data-independent acquisition (DIA) method with a 120K resolution Orbitrap MS1 scan and 12 m/z isolation window, 30K resolution Orbitrap MS2 scans for precursors from 400-1000 m/z. Data .raw files were converted to .mzML using MSConvert 3.0.21251-d2724a5 and spectral libraries built using MSFragger-DIA (PMID: 28394336) (w/FragPipe version 22.0 and MSFragger version 4.1) with quantification through DIA-NN versions 1.8.2 (PMID: 31768060). The database search was against the UniProt human database (downloaded 2024-01-24) with supplemental spike-in of common contaminants, containing 20477 sequences and 20477 reverse-sequence decoys. For the MSFragger analysis, both precursor and (initial) fragment mass tolerances were set to 20 ppm. Spectrum deisotoping, mass calibration, and parameter optimization were enabled. Enzyme specificity was set to “stricttrypsin” and up to two missed trypsin cleavages were allowed. Oxidation of methionine, acetylation of protein N-termini, −18.0106 Da on N-terminal Glutamic acid, and −17.0265 Da on N-terminal Glutamine and Cysteine were set as variable modifications. Carbamidomethylation of Cysteine was set as a fixed modification. Maximum number of variable modifications per peptide was set to 3. FragPipe/DIA-NN output files were processed and analyzed using the Perseus software package v2.0.11.0. Expression columns (protein MS intensities) were log2 transformed and normalized by subtracting the median log2 expression value from each expression value within each MS run. Missing values were imputed from a normal distribution; a width of 2 units and a downshift of 1.8 units was applied. For statistical testing of significant differences in expression, a two-sample Student’s t test was applied. A permutation based false discovery rate of 0.05 was used to correct for multiple comparison.

### Single Cell RNA Sequencing

Organoids at week 14 of differentiation were used for single-cell RNA sequencing (scRNA-seq). For each genotype, 5-6 organoids from each of two independent clones were pooled, yielding four multiplexed samples (2 genotypes × 2 clones). The organoids pooled within each clonal sample were derived from at least two independent rounds of differentiation. Organoids were dissociated into single-cell suspensions using the Papain Dissociation System (Worthington Biochemical Corporation). Cells were processed with the Chromium GEM-X Single Cell 3′ v4 assay (10x Genomics) according to the manufacturer’s protocol. Complementary DNA quality was assessed using an Agilent TapeStation 4200, and libraries were sequenced on an Illumina NextSeq 2000.

### Processing and Analysis of Single-Cell Sequencing Data

FASTQ files were aligned to a modified version of the human genome (GRCh38) using CellRanger 6.1.2 (10x Genomics). Cell Ranger gene expression matrix outputs were then loaded into Seurat 5.5. Data from the samples were converted to a Seurat object and metadata regarding genotype and clone was added to the object. The object was subset, extracting cells in which >800 and <7500 RNA features were expressed and <5% of the genes expressed were mitochondrial. Specifically, cells were retained if they met all of the following criteria: total UMI counts between 1,000 and 40,000; number of detected genes between 800 and 7,500; mitochondrial read percentage below 5%; ribosomal read percentage below 40%; and a log10-transformed genes-per-UMI ratio greater than 0.80. Principal Component Analysis (PCA) was then run on the post-filtered cells, followed by UMAP dimensionality reduction using the first 20 PCs. Shared nearest neighbors for each cell and cluster were identified. Cell-type labels were transferred from a downsampled (≤500 cells per subtype) developing human cortex scRNA-seq reference atlas ^38^(Nano et al 2025) to our query dataset using Seurat’s anchor-based label transfer. Transcriptional concordance between query clusters and reference cell types was assessed by Pearson correlation of pseudobulk average expression profiles across shared variable genes, visualized as a hierarchically clustered heatmap (Ward’s method). Cells annotated as interneurons (IN) were subset from the full Seurat object for independent re-analysis. Gene expression was log-normalized and the top 2,000 highly variable genes were identified using the variance-stabilizing transformation method. Data were scaled and principal component analysis was performed. Cells were clustered using the Louvain algorithm and were visualized by UMAP. For pseudobulk gene expression analysis, raw UMI counts were aggregated across cells sharing the same cell type, genotype, and clone, yielding one pseudobulk profile per combination (2 clones per genotype). Differential expression between TAOK1 ^S111F het^ and Ctrl^wt^ was tested separately within each of the annotated cell types using DESeq2 (v[1.50.2]; Wald test, Benjamini–Hochberg-adjusted p-values), with dispersion estimated across the full transcriptome before subsetting results to genes of interest. For YAP-TEAD target gene set analysis, Seurat’s AddModuleScore() function was used to compute a relative enrichment score for the YAP-TEAD gene set. Enrichment score is average expression of the signature genes) − (average expression of the sampled control genes. Statistical significance was assessed using a Wilcoxon rank-sum test comparing the distribution of per-cell module scores between TAOK1 ^S111F het^ and Ctrl^wt^ cells, grouped by cluster. Pseudotime trajectory analysis was performed using Monocle 3 package. The Seurat object was converted to a Monocle 3 cell_data_set object and cells were subset by genotype so that wild-type and heterozygous cells were analyzed separately. For each genotype, cells were clustered with Monocle 3’s cluster cells function, and the resulting partitions and the previously defined Seurat clusters were visualized on the UMAP embedding carried over from the Seurat workflow. A principal graph was fit to the data with the partition restriction disabled so that a single trajectory was learned across all cells rather than separately within each partition. Cells were then ordered along the trajectory, with the root placed in the radial glial cell cluster, and pseudotime values were visualized on the UMAP embedding.

### Microscopy and Image Analysis

All Imaging was performed on a Nikon Ti2 Eclipse-CSU-X1 confocal spinning disk microscope equipped with four laser lines (405, 488, 561, and 670 nm) and an sCMOS Andor camera for image acquisition with the exception of image 1g. The microscope was caged within the OkoLab environmental control setup enabling temperature and CO_2_ control during live imaging. Imaging was performed using Nikon 1.49 100× Apo, 60× or 40× oil objective lenses. Figure 1g was captured on the Zeiss Elyra7 Lattice SIM2 structural illumination microscope. Imaging was performed on a Zeiss Axio Observer 7 - Lattice 648 SIM inverted microscope for super resolution microscopy equipped with four laser lines – 405 649 nm (50mW), 488 nm (100mW), 561 nm (100mW) and 642 nm (150mW) – and the ZEN imaging 650 software. Two Hamamatsu ORCA-Fusion BT sCMOS cameras were used for simultaneous 651 image acquisition. Imaging was performed using Plan-Apochromat 25x/1.40 oil DIC objective. Tiles were stitched using the stitching function available in the Zeiss software. All image analyses and quantification of fluorescence intensity was performed using Fiji ImageJ. Ki-67-, Cyclin A2-, and DAPI-positive cells in NPCs were counted manually by two blinded scorers and one unblinded scorer, with each round of differentiation counted by a different scorer. In organoids, Ki-67/Cyclin A2/DAPI positive cells were quantified using CellProfiler (v4.2.8), which identified primary objects in each channel using the IdentifyPrimaryObjects module with a nuclear diameter range of 40-120 pixels. Marker-positive counts were expressed as a percentage of DAPI-positive cells. Colocalization of DAPI and YAP1 was also quantified in CellProfiler, and Pearson’s correlation coefficient was reported.

### Real-time quantitative polymerase chain reaction (RT-qPCR)

Total RNA was isolated using the Monarch® Spin RNA Isolation Kit (New England Biolabs, USA) and template complementary DNA was prepared by reverse transcription using the LunaScript RT Supermix ((New England Biolabs, USA). Quantitative analysis of relative gene expression was determined via CFX-96 Real Time PCR detection system (Bio-Rad, UK). Primers and sequences are added in supplementary table.

### Statistics

Statistics were performed in GraphPad software Prism 10.0. Multiple groups were analyzed using ANOVA, while two group comparisons were made using unpaired t test unless otherwise stated. Statistically *p* value less than 0.05 was considered significant. All experiments were done in triplicate, and experimental sample size and *p* values are indicated with the corresponding figures.

---

**Extended Data Figure 1.**
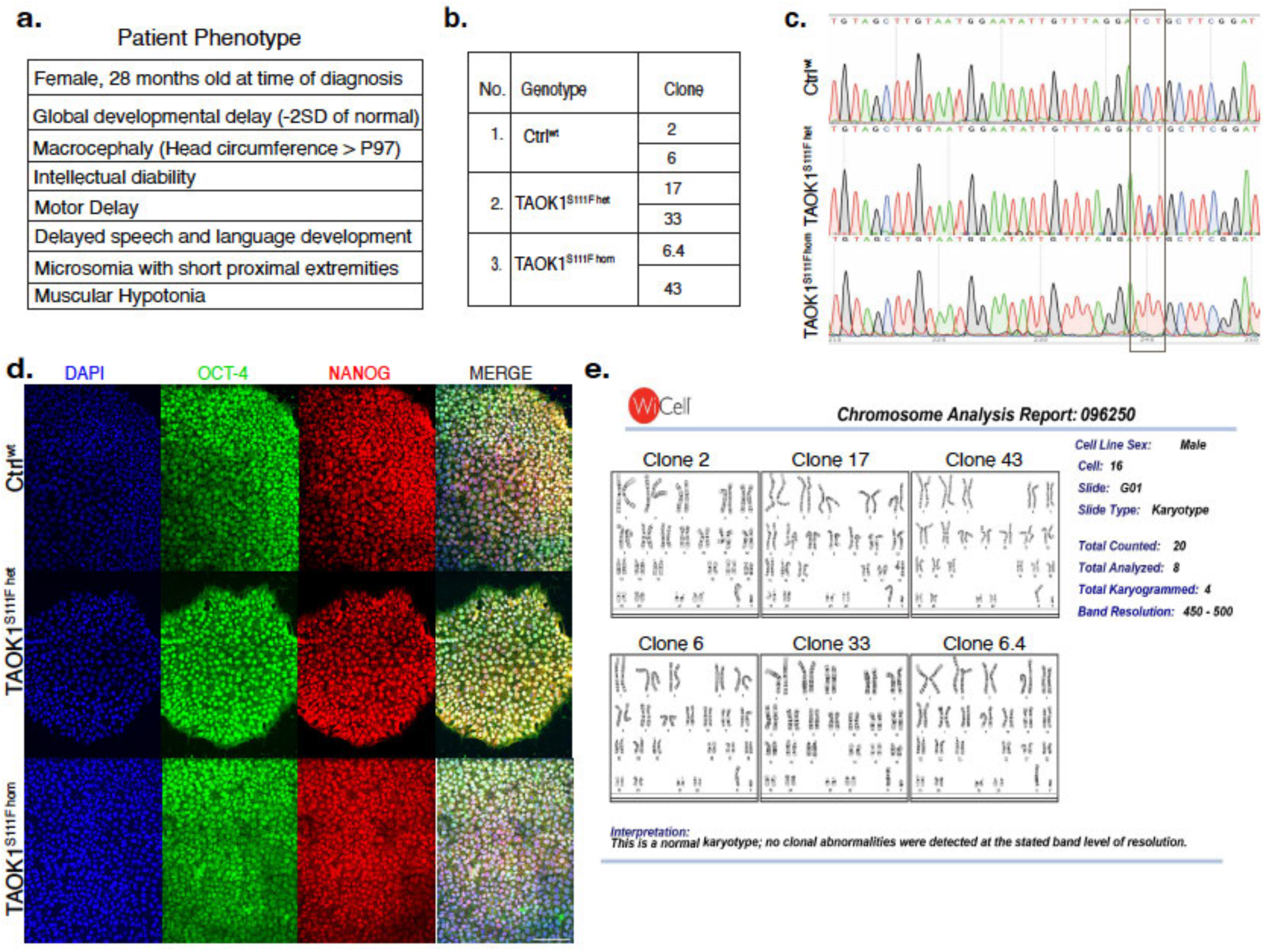
Generation of iPSC lines harboring S111F pathogenic TAOK1 variant. **(a)** Clinical features of patient carrying a heterozygous TAOK1 S111F mutation reported by Dulovic-Mahlow et al., 2019. **(b)** Clones of engineered iPSCs (WTC11) with heterozygous and homozygous S111F mutations along with isogenic controls used in this study. **(c)** Sequencing traces show the targeted TAOK1 exon 5 and resulting edit for the S111 to F substitution (TCT→TTT) in heterozygous and homozygous lines shown alongside the isogenic control. **(d)** Representative immunofluorescent images of Ctrl^WT^, TAOK1^S111F het^ and TAOK1^S111F hom^ iPSCs immunostained with DAPI (blue), OCT-4 (green), and NANOG (red) confirming pluripotent identity. **(e)** Karyotype analysis of all genome-edited iPSC clones confirming absence of any large chromosomal aberrations.

**Extended Data Figure 2.**
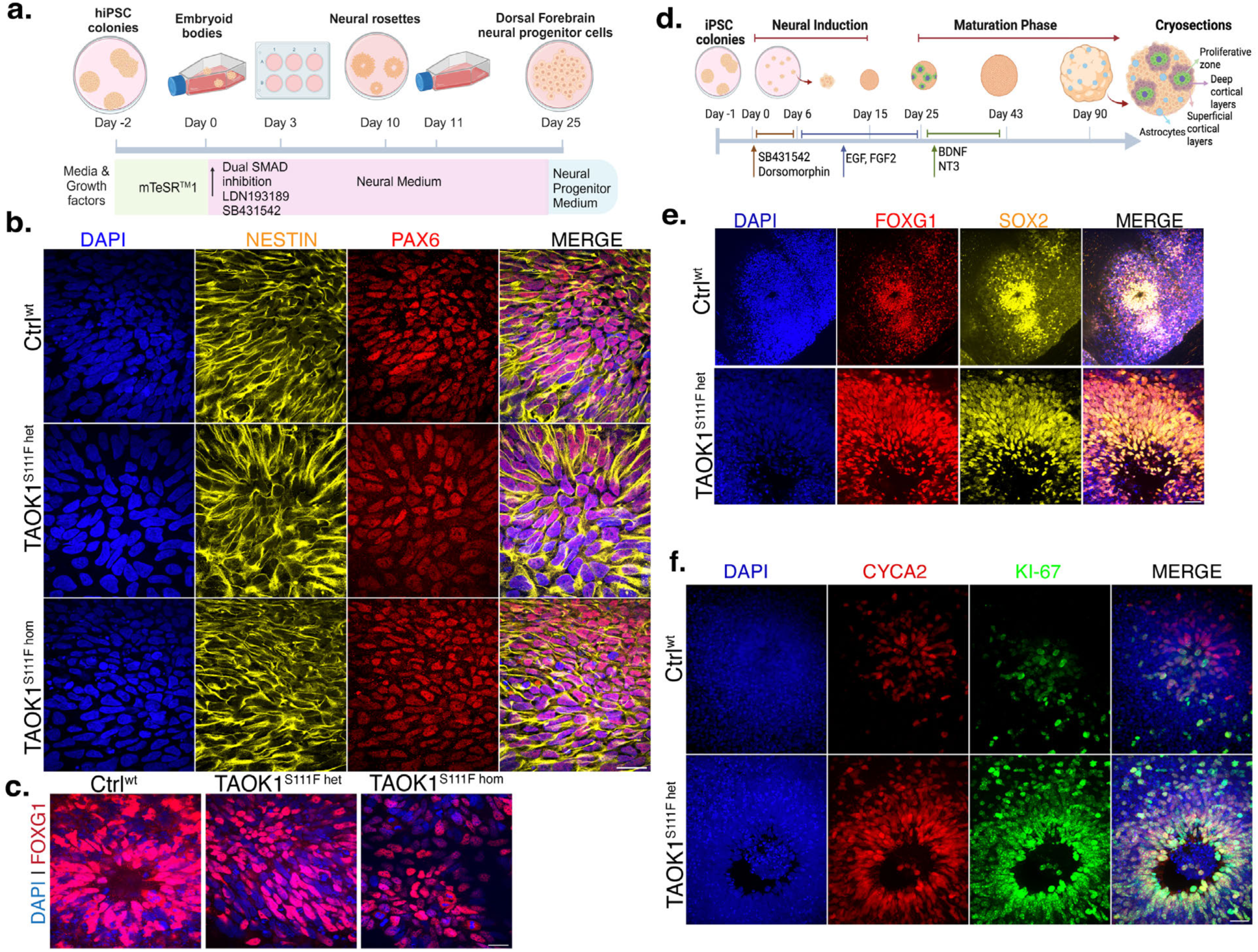
Dorsal forebrain neural progenitors and cortical organoids differentiated from iPSC lines. **(a)** Schematic shows the workflow for differentiation of iPSCs into dorsal forebrain neural progenitors using dual SMAD inhibition. **(b)** Representative immunofluorescent images of Ctrl^WT^, TAOK1^S111F het^, and TAOK1^S111F hom^ NPCs immunostained for Nestin (yellow), PAX6 (red) and DAPI (blue) confirming dorsal neuroprogenitor identity. Scalebar, 10µm. **(c)** Representative immunofluorescent images of Ctrl^WT^, TAOK1^S111F het^ and TAOK1^S111F hom^ NPCs immunostained for FOXG1 (red) and DAPI (blue) confirming dorsal forebrain identity. Scalebar, 10µm. **(d)** Schematic depicts workflow for generation of dorsal forebrain cortical organoids from iPSCs. **(e)** Representative cortical organoid sections of Ctrl^wt^, TAOK1^S111F het^ and TAOK1^S111F hom^ immunostained for FOXG1 (red), SOX2 (yellow), and DAPI (blue) confirming dorsal forebrain organoid identity. Scalebar, 10µm. **(f)** Representative immunofluorescent images of 10-week Ctrl^WT^, TAOK1^S111F het^, and TAOK1^S111F hom^ NPCs cortical organoid sections stained for Ki67 (green), Cyclin A2 (red), and DAPI (blue). Scalebar, 10µm.

**Extended Data Figure 3:**
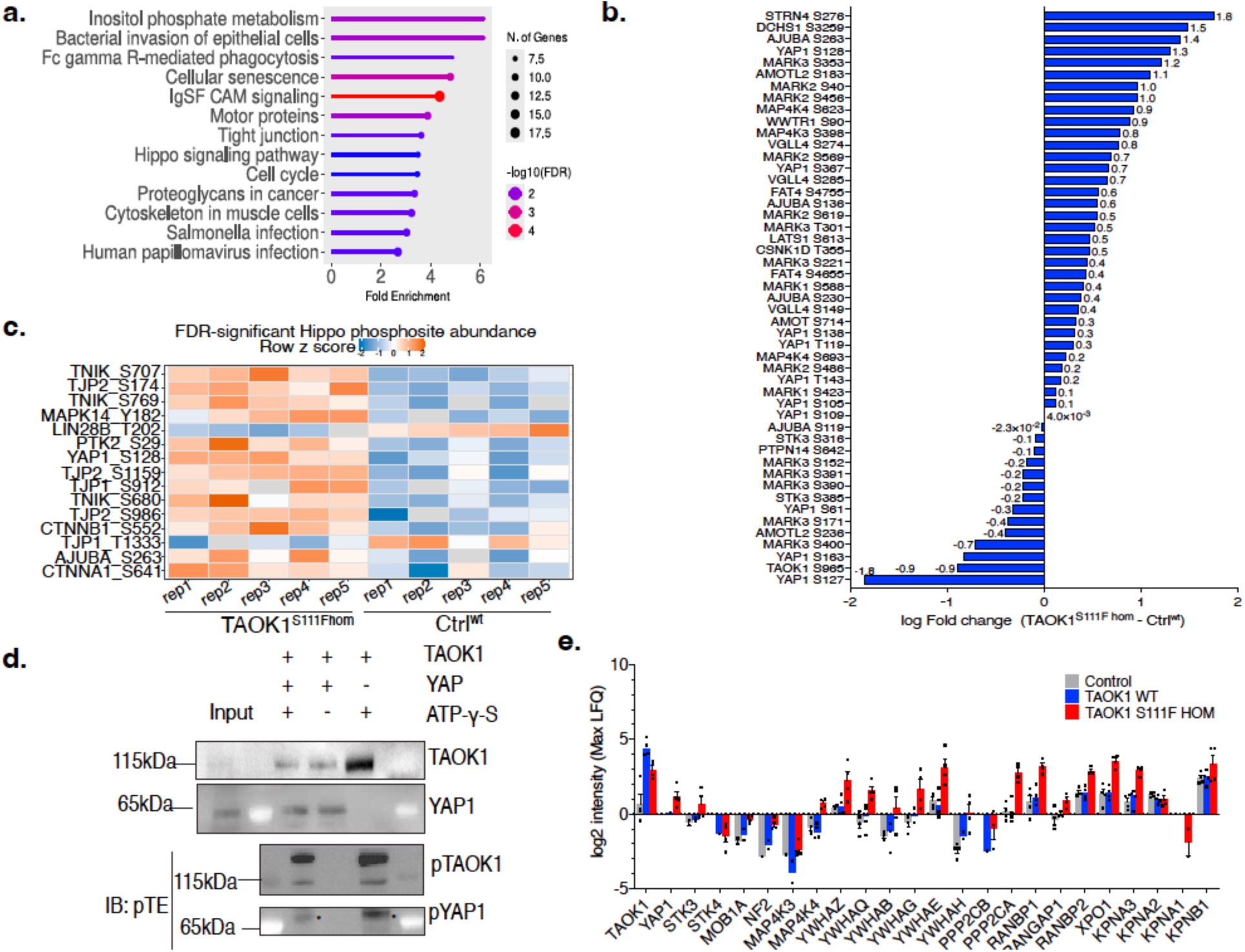
TAOK1 interacts with YAP1 but does not phosphorylate YAP1. **(a)** Lollipop plot of the top enriched KEGG pathways from phosphoproteomic profiling of Ctrl^WT^ and TAOK1^S111F^ NPCs. The x-axis shows fold enrichment, point color denotes −log_10_(FDR), and point size corresponds to the number of associated genes. **(b)** Bar chart of fold-change in phosphorylation of all known Hippo pathway components identified in our phosphoproteomics data comparing Ctrl^WT^ and TAOK1^S111F hom^ NPCs. Differential abundance was assessed by limma and log fold change for each protein is indicated on top of individual bars. **(c)** Heat map showing all FDR significant Hippo phosphosite abundance between the Ctrl^WT^ and TAOK1^S111F hom^ NPC. **(d)** Endogenous YAP1 was immunoprecipitated from Ctrl^WT^ NPC lysates and incubated with purified TAOK1 and ATPγS in an in vitro kinase reaction. Presence of thiophosphorylated proteins was detected using an antibody against thiophosphate ester. At molecular weight 115KDa, pTAOK1 was detected however no phosphorylated YAP1 was detected at 65KDa. Asterisk indicates a non-specific background band above 65KDa which is also seen in condition with no YAP1 protein. **(e)** Bar plot of log_2_ LFQ (label free quantification) intensity for key Hippo pathway regulators YAP/TAZ, 14-3-3 proteins, phosphatases, and nuclear import/export proteins in Ctrl^WT^ and TAOK1^S111F hom^ NPC lysates immunoprecipitated with TAOK1 antibody.

**Extended Data Figure 4:**
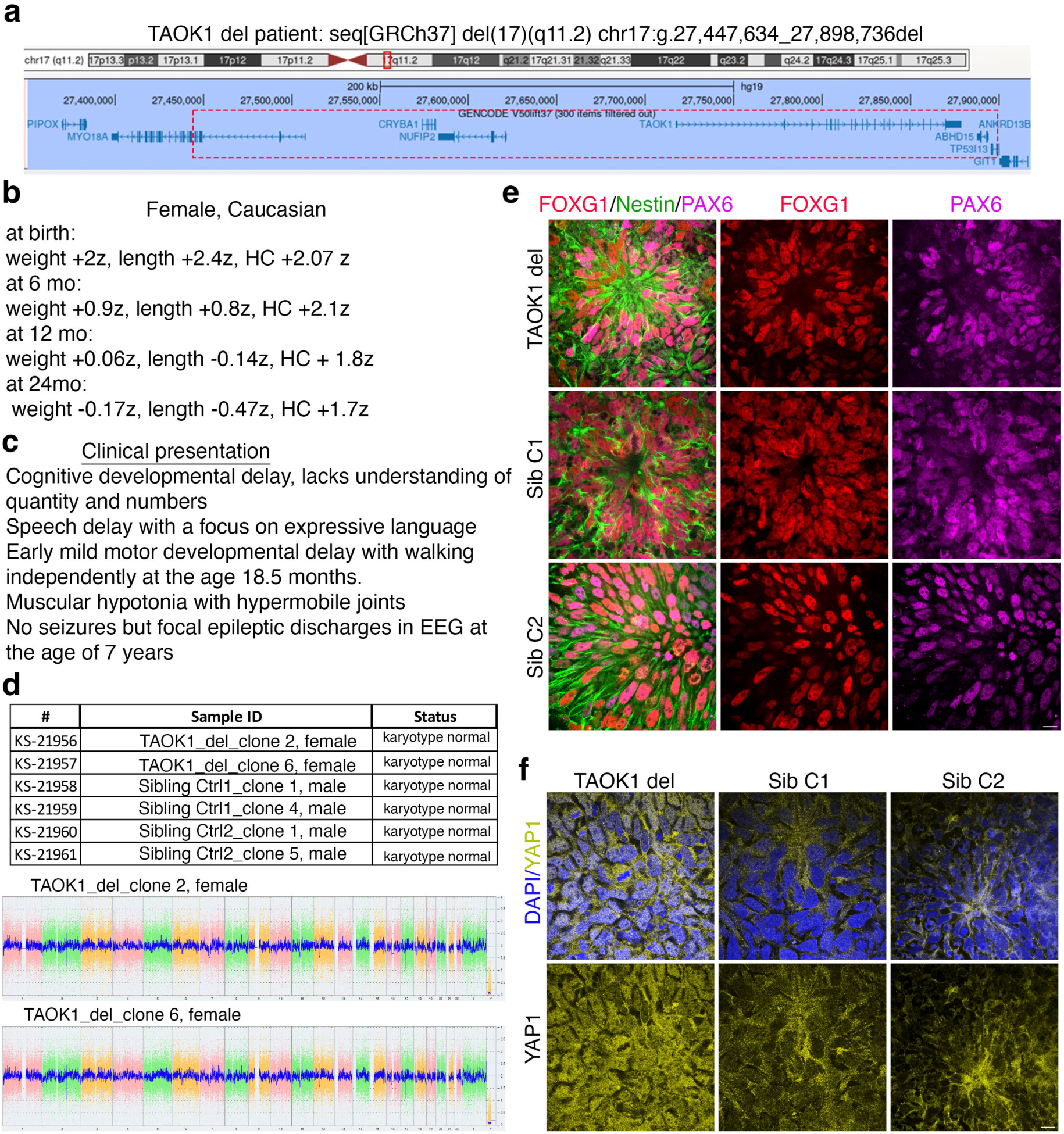
Characterization of TAOK1 deletion patient-derived neural progenitors. **(a)** Schematic of chromosome 17 depicting the coordinates of the genomic deletion in the TAOK1 deletion patient. Deletion includes TAOK1 and 4 other genes, none of which have a known association with neurodevelopmental deficit or macrocephaly. **(b)** Growth trajectory of head circumference, weight and length of the patient from birth up to 24 months. **(c)** Main clinical presentation of the TAOK1 deletion patient up until the last evaluation at 7yr of age. **(d)** Representative immunofluorescent images of NPCs differentiated from the patient and unaffected sibling iPSCs immunostained for Nestin (green), PAX6 (magenta) and FOXG1(red). Scale bar, 10µm. **(e)** Karyotype analysis of patient and unaffected siblings iPSC clones, confirming absence of any large chromosomal aberrations. **(f)** Representative immunofluorescence images of NPCs differentiated from patient iPSCs carrying a heterozygous TAOK1 deletion, immunostained for YAP1 (yellow) and DAPI (blue). NPCs differentiated from unaffected siblings iPSCs serve as controls. Scale bar, 10µm.

**Extended Data Figure 5:**
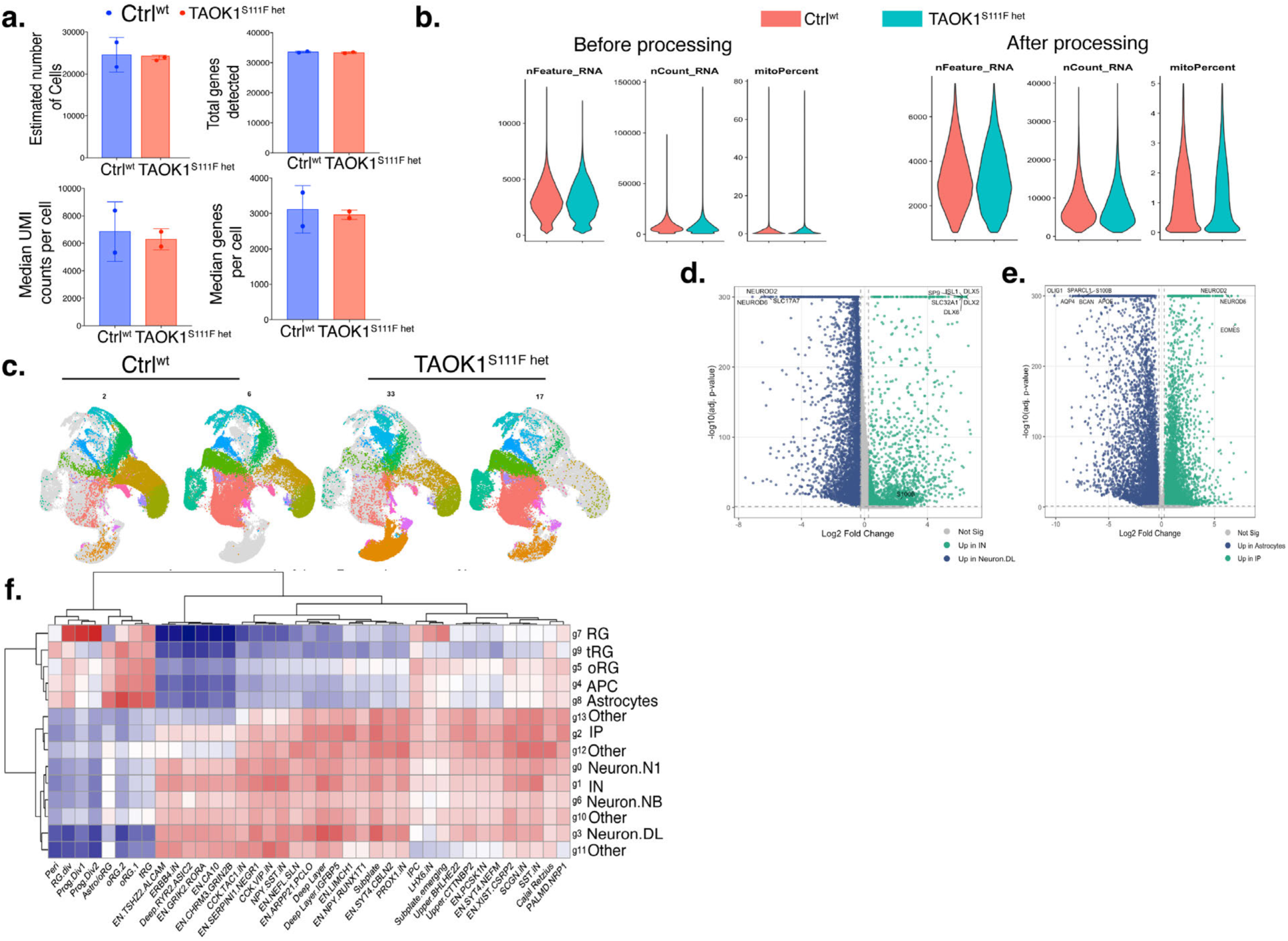
Quality control metrics and cluster composition of scRNA-seq datasets. **(a)** Bar graphs plot estimated number of cells, total genes detected, median UMI counts per cell, and median genes per cell between the Ctrl^wt^ and TAOK1^S111F het^ cortical organoids scRNA SEQ run. **(b)** Violin plots show number of genes, number of RNA counts, and percent mitochondrial genes before and after processing the scRNA seq data set. **(c)** UMAP plots of scRNA-seq data from Ctrl^wt^ and TAOK1^S111F het^ cortical organoids with cells grouped by unbiased clustering and clones for each genotype. **(d)** Volcano plot of differentially expressed genes between interneuron (IN) and deep-layer excitatory neuron (Neuron.DL) clusters, pooled across genotypes. The x-axis shows log_2_ fold change and the y-axis shows −log_10_(adjusted p-value). Differential expression was assessed using the Wilcoxon rank-sum test (Seurat FindMarkers), with p-values adjusted for multiple testing by Bonferroni correction. Dashed vertical lines indicate the log2 fold-change threshold (±0.25); the dashed horizontal line indicates the adjusted p-value significance threshold (0.05). Genes enriched in IN are shown in teal; genes enriched in Neuron.DL are shown in blue; non-significant genes are shown in gray. Selected interneuron and excitatory neuron marker genes are labeled. **(e)** Volcano plot of differentially expressed genes between astrocytes and intermediate progenitor cells (IP) clusters, pooled across genotypes. The x-axis shows log_2_ fold change and the y-axis shows −log_10_(adjusted p-value). Differential expression was assessed using the Wilcoxon rank-sum test (Seurat FindMarkers), with p-values adjusted for multiple testing by Bonferroni correction. Dashed vertical lines indicate the log_2_ fold-change threshold (±0.25); the dashed horizontal line indicates the adjusted p-value significance threshold (0.05). Genes enriched in IP are shown in teal; genes enriched in astrocytes are shown in blue; non-significant genes are shown in gray. Selected IP and astrocyte marker genes are labeled. **(f)** Heatmap depicts pairwise transcriptional correlation between cell clusters identified by scRNA seq of week 14 cortical organoids (rows) and cell types annotated in the developing human cortex scRNA-seq meta-atlas (*Nano et al., 2025,* columns). Correlations were calculated as Pearson coefficients on pseudobulk average expression profiles. Rows and columns are labeled by principal cell-type/cluster identity; dendrograms show hierarchical clustering (Ward’s method) of correlation distances.

**Extended Data Figure 6:**
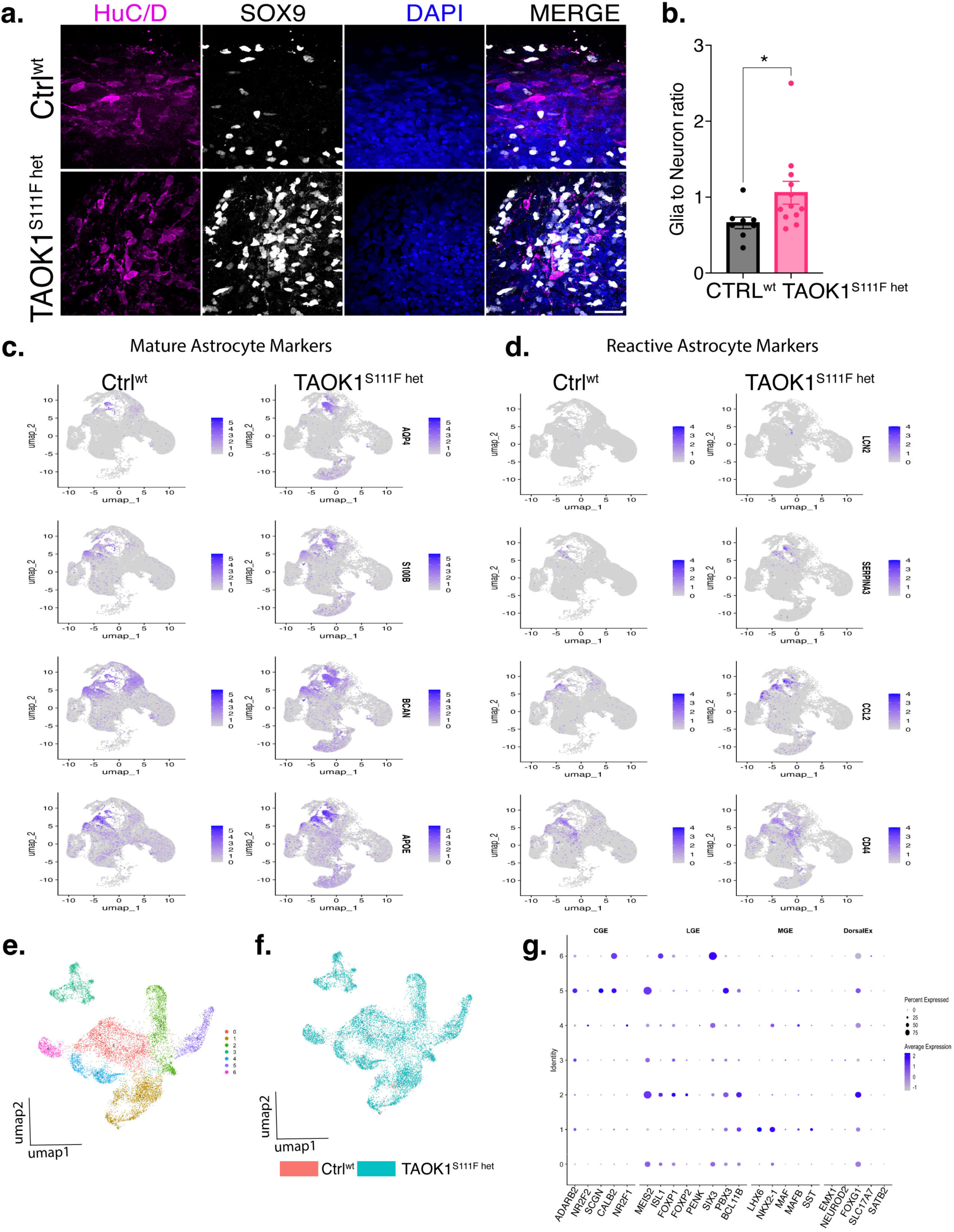
Transcriptional analyses of the astrocyte and inhibitory neuron population in cortical organoids. **(a)** Representative immunofluorescence images of SOX9 and HuC/D staining in Ctrl^WT^ and TAOK1^S111F het^ dorsal forebrain cortical organoid (DIV100) slices. Scale bar: 25µm. **(b)** Bar plot shows (mean ± SEM) ratio of (SOX9+ive) glia to (HuC/D+ive) neuronal cells per ROI in Ctrl^WT^ and TAOK1^S111F het^ cortical organoid sections at DIV 100 (n=6 organoids per genotype). Statistical significance was assessed by student’s t test (Mann–Whitney test), *p< 0.05. **(c)** Feature plots of mature astrocyte markers in Ctrl^WT^ and TAOK1^S111F het^ cells. **(d)** Feature plots of reactive astrocyte markers in Ctrl^wt^ and TAOK1^S111F het^ cells. **(e)** UMAP visualization of inhibitory neuronal (IN) cells after subclustering, colored by unsupervised cluster identity, and **(f)** by genotype. **(g)** Dot plot shows expression of canonical marker genes for CGE, LGE, MGE derived IN and dorsal excitatory neuronal populations. Color intensity reflects expression level (dark blue, high expression); dot size reflects the percentage of cells expressing each marker.

**Extended Data Figure 7:**
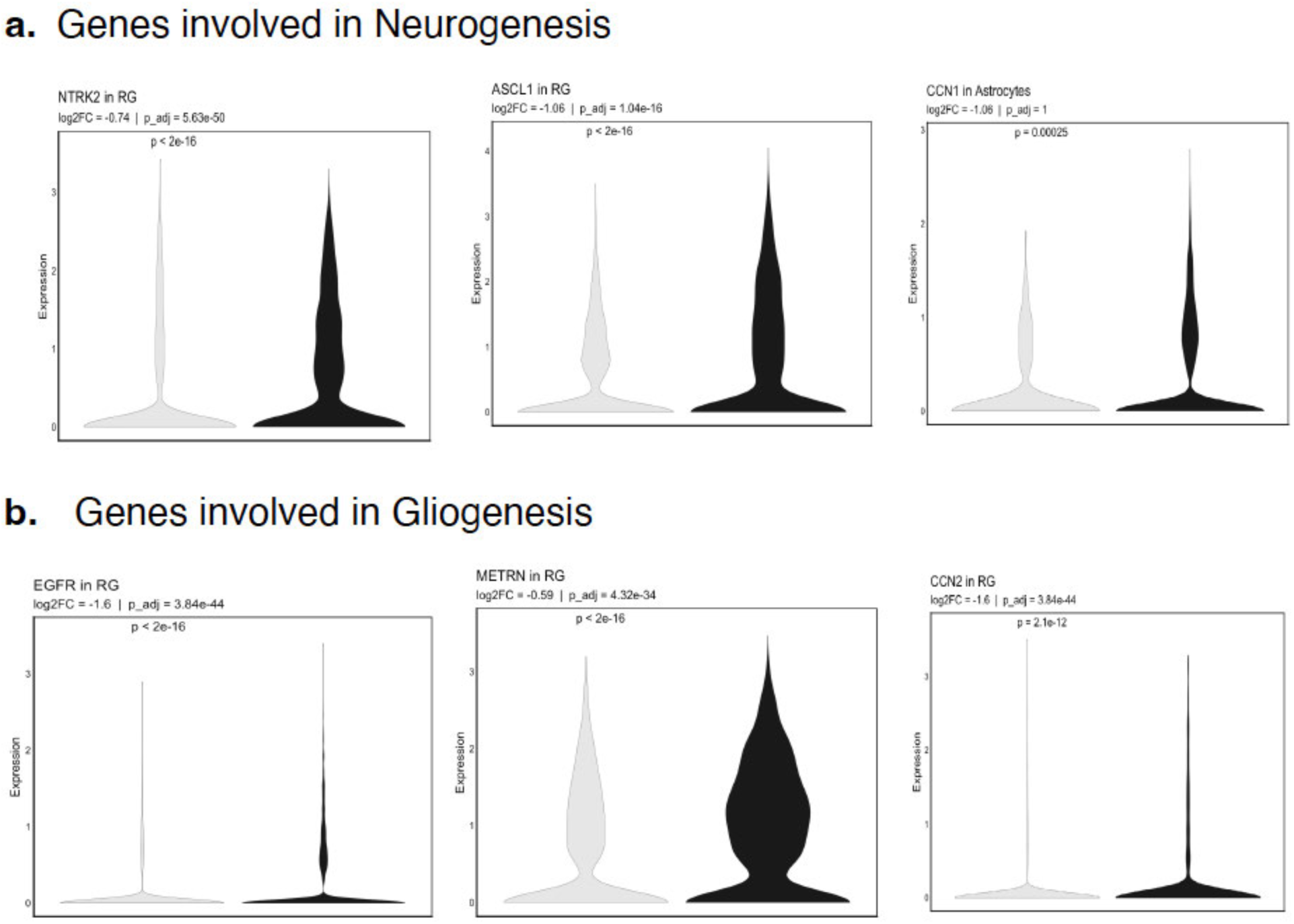
YAP-TEAD target gene expression analysis. **(a)** Violin plots show distribution of expression of select YAP-TEAD genes that promote neurogenesis, *NTRK2, ASCL1*, and *CCN1* between Ctrl^wt^ and TAOK1^S111F het^ in the indicated cell cluster. **(b)** Violin plots show distribution of expression of select YAP-TEAD genes involved in gliogenesis, *EGFR, METRN*, and *CCN2* between Ctrl^wt^ and TAOK1^S111F het^ in the indicated cell population.

